# Action potential-coupled Rho GTPase signaling drives presynaptic plasticity

**DOI:** 10.1101/2020.10.07.330126

**Authors:** Shataakshi Dube, Bence Rácz, Walter E. Brown, Yudong Gao, Erik J. Soderblom, Ryohei Yasuda, Scott H. Soderling

## Abstract

In contrast to their postsynaptic counterparts, the contributions of activity-dependent cytoskeletal signaling to presynaptic plasticity remain controversial and poorly understood. To identify and evaluate these signaling pathways, we conducted a proteomic analysis of the presynaptic cytomatrix using *in vivo* biotin identification (iBioID). The resultant proteome was heavily enriched for actin cytoskeleton regulators, including Rac1, a Rho GTPase that activates the Arp2/3 complex to nucleate branched actin filaments. Strikingly, we find Rac1 and Arp2/3 are closely associated with presynaptic vesicle membranes and negatively regulate synaptic vesicle replenishment at both excitatory and inhibitory synapses. Using optogenetics and fluorescence lifetime imaging, we show this pathway bidirectionally sculpts short-term synaptic depression and that its presynaptic activation is coupled to action potentials by voltage-gated calcium influx. Thus, this study provides a new proteomic framework for understanding presynaptic physiology and uncovers a previously unrecognized mechanism of actin-regulated short-term presynaptic plasticity that is conserved across cell types.

## INTRODUCTION

Dynamic tuning of neurotransmitter release in response to patterns of activity is a fundamental process that ultimately governs how experience modulates neural networks. During bursts of high-frequency firing, the complex interplay between presynaptic calcium levels and vesicle availability can result in a transient enhancement or reduction of synaptic strength, a process known as short-term synaptic plasticity (Regehr, 2012). Although recent work has clarified some of the calcium sensors important for short-term enhancement during facilitation (Jackman and Regehr, 2017; Jackman et al., 2016) and augmentation (Xue et al., 2018), the signaling molecules that sense action potentials to translate other forms of short-term plasticity are still poorly understood. Reduction of release during short-term depression (STD) is generally thought to reflect the depletion of the readily releasable pool of synaptic vesicles. However, at many synapses, vesicle depletion cannot fully account for the extent of depression (Bellingham and Walmsley, 1999; Byrne, 1982; Chen et al., 2004; Hsu et al., 1996; Kraushaar and Jonas, 2000; Parker, 1995; Sullivan, 2007; Thomson and Bannister, 1999; Waldeck et al., 2000; Xu and Wu, 2005; Zucker and Bruner, 1977), suggesting the presence of additional unknown activity-dependent signaling mechanisms driving STD.

The actin cytoskeleton has long been implicated in many stages of the synaptic vesicle cycle that could modulate short-term plasticity, including exocytosis, endocytosis, vesicle trafficking, and reserve pool clustering (Cingolani and Goda, 2008; Rust and Maritzen, 2015). Yet these potential roles have been controversial, as actin depolymerizing agents have enhanced, reduced, or had no effect on each of these processes depending on the study (Cole et al., 2000; Darcy et al., 2006; Gaffield et al., 2006; Gramlich and Klyachko, 2017; Lee et al., 2012; Morales et al., 2000; Sakaba and Neher, 2003; Sankaranarayanan et al., 2003). These pharmacological manipulations, while powerful, may not be the ideal method to reveal the diverse functions and regulation of presynaptic actin, because they influence the entire actin cytoskeleton. They do not specifically probe the unique actin pools that exist within different subcellular compartments (Papandreou and Leterrier, 2018). Indeed, many aspects of postsynaptic physiology have been clarified by genetic analyses of actin signaling cascades within dendritic spines. These studies have revealed that distinct pools of actin sculpt dendritic spine morphology, modulate adhesion, and regulate plasticity mechanisms such as the anchoring and trafficking of glutamate receptors (Spence and Soderling, 2015). These different pools are tightly regulated by the Rho-family GTPases (including RhoA, Rac1, and Cdc42), which act on effector proteins to control actin filament assembly and disassembly during both baseline transmission and synaptic plasticity (Hedrick and Yasuda, 2017; Murakoshi et al., 2011; Tolias et al., 2011). Furthermore, these signaling pathways are heavily implicated in neurological diseases such as intellectual disability, autism, and schizophrenia (Spence and Soderling, 2015; Yan et al., 2016), highlighting the importance of synaptic actin for proper neural function. Given the clear links between actin turnover and postsynaptic plasticity, it is therefore surprising that there is little evidence supporting a role for the presynaptic actin cytoskeleton or its signaling molecules in mechanisms of short-term presynaptic plasticity. Some studies have even suggested that presynaptic actin remodeling is only important during synapse maturation (Shen et al., 2006; Yao et al., 2006).

Here, we uncover a new, conserved role for Rho-family GTPase signaling in driving STD at both glutamatergic and GABAergic presynaptic terminals. First, in order to enable genetic analysis of the presynaptic cytoskeleton, we defined the actin signaling pathways present in presynaptic terminals. These proteins have not been systematically identified because the presynaptic cytomatrix cannot be biochemically purified, limiting previous studies of the presynaptic proteome to synaptic vesicles and the active zone. To capture a larger fraction of the presynaptic cytomatrix, we used *in vivo* Biotin Identification (iBioID) and localized the promiscuous biotin ligase BioID2 to presynaptic terminals by fusing it to Synapsin, a presynaptic actin-binding protein (Doussau and Augustine, 2000; Greengard et al., 1994). Similar to our previous work isolating the proteomes of inhibitory postsynapses (Uezu et al., 2016) and dendritic filopodia (Spence et al., 2019), this approach led to the mass spectrometry-based identification of 200 proteins within presynaptic terminals of the hippocampus and cortex. This network of presynaptic proteins was highly enriched for regulators of the actin cytoskeleton and converged on a Rac1-Arp2/3 signaling pathway that leads to the *de novo* nucleation of branched actin filaments (Higgs and Pollard, 2001; Mullins et al., 1998). While Rac1 and Arp2/3 have established roles at the postsynapse (Hedrick and Yasuda, 2017; Kim et al., 2013; Spence et al., 2016; Tolias et al., 2011), here we discovered that Rac1 and Arp2/3 are also closely associated with presynaptic vesicle membranes *in vivo*. We developed genetic, optogenetic, and electrophysiological strategies to specifically isolate presynaptic effects and demonstrated that Rac1-Arp2/3 signaling negatively regulates synaptic vesicle replenishment and can bidirectionally alter STD. By imaging a Rac1 activity sensor (Hedrick et al., 2016) in presynaptic terminals, we also found that Rac1 activation is coupled to action potential trains via voltage-gated calcium influx. Thus, Rac1 and branched actin have an important, previously uncharacterized presynaptic role in sculpting short-term synaptic plasticity. These results define a new activity-dependent signaling mechanism that contributes to STD and is conserved across cell types. This also challenges the prevailing view that the Rac1-Arp2/3 pathway functions largely at excitatory postsynapses, prompting re-evaluation of its mechanism in neurodevelopmental disorders.

## RESULTS

### Identification of the proteomic composition of the presynaptic cytomatrix *in vivo*

Current knowledge about presynaptic actin regulation at mature synapses is limited to the discovery of both pre- and post-synaptic effects in a few genetic knockout studies (Connert et al., 2006; Wolf et al., 2015; Xiao et al., 2016). A larger inventory of presynaptic actin regulators is still lacking due to the inability of traditional biochemical methods to isolate the presynaptic cytomatrix, where actin signaling likely occurs. Proteomic studies from isolated synaptic vesicles and active zone fractions, although powerful, have identified few actin signaling molecules (Abul-Husn et al., 2009; Boyken et al., 2013; Burre et al., 2006; Coughenour et al., 2004; Morciano et al., 2009; Morciano et al., 2005; Takamori et al., 2006; Weingarten et al., 2014; Wilhelm et al., 2014), despite actin being the most abundant cytoskeletal element in presynaptic terminals (Wilhelm et al., 2014).

We turned to a proximity-based proteomics approach, *in vivo* Biotin Identification (iBioID), in which the promiscuous biotin ligase BioID2 is fused to a protein in a compartment of interest, and nearby biotinylated proteins are identified by mass spectrometry (Kim et al., 2016; Spence et al., 2019; Uezu et al., 2016). To direct BioID2’s activity towards the presynaptic cytomatrix, we created a Synapsin1a fusion protein with a flexible 4x[GGGGS] linker (Figure 1A). Synapsin is a synaptic vesicle protein that is also known to bind actin (Doussau and Augustine, 2000; Greengard et al., 1994), making it the ideal bait for discovering presynaptic actin signaling pathways. Importantly, Synapsin has been tagged previously with GFP without disrupting its presynaptic targeting (Gitler et al., 2004b). To validate this approach, we expressed BioID2-Synapsin, untargeted BioID2, and GFP in cultured hippocampal neurons and incubated them with exogenous biotin (Figure 1—figure supplement 1A-C). BioID2-Synapsin was enriched in presynaptic boutons similarly to Bassoon, an active zone marker, while the localization of BioID2 was indistinguishable from GFP, confirming it acts as a soluble fill (Figure 1—figure supplement 1D). The biotinylation activity of BioID2-Synapsin was also significantly enhanced in presynaptic terminals in comparison to BioID2 alone (Figure 1—figure supplement 1E).

**Figure 1.**
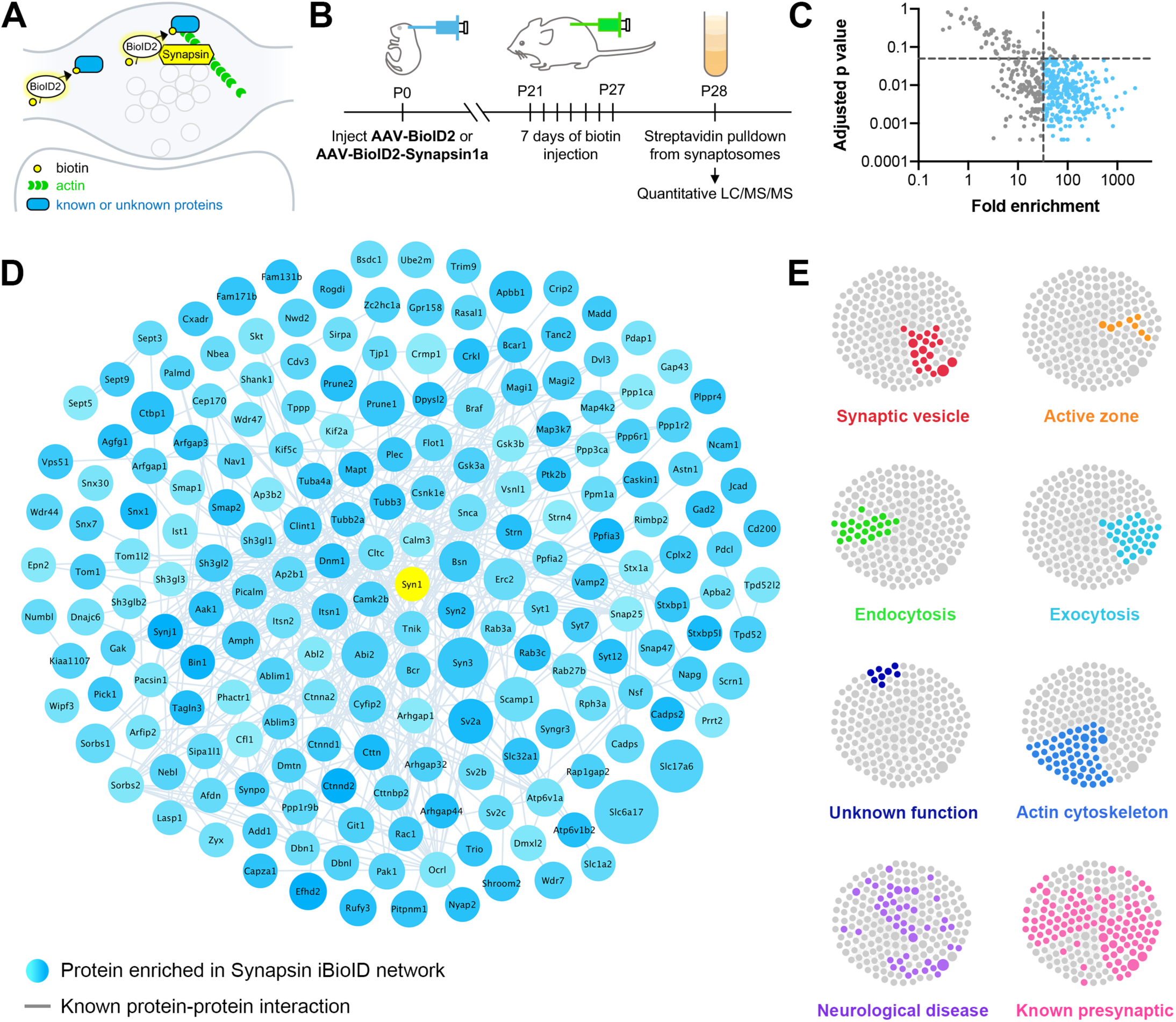
Identification of the proteomic composition of the presynaptic cytomatrix using *in vivo BioID.* **(A)** Schematic of the iBioID approach in presynaptic terminals. **(B)** Timeline of *in vivo* injections and sample collection. **(C)** Filters used to select proteins based on fold enrichment over negative control and FDR adjusted p-value (t-tests). **(D)** Synapsin iBioID identified a rich network of 200 known and previously unknown proteins enriched in presynaptic terminals. Node titles correspond to gene name, size represents fold enrichment over the BioID2 negative control (range 32.7 - 2275.1), shading represents FDR adjusted p-value with light blue being a lower p-value and darker blue a higher p-value (range 0.0003 - 0.049). Edges are previously reported protein-protein interactions in the HitPredict database or by hand annotation. **(E)** Clustergrams of proteins that are in synaptic vesicles (red, n=20/200 proteins) or active zones (orange, n=8); involved in endocytosis (green, n=22), exocytosis (cyan, n=30), or actin regulation (blue, n=54); have unknown function (navy, n=8); are implicated in neurological diseases (purple, n=46) as identified through DAVID analysis or hand annotation; and are known to be presynaptic (pink, n=108).

With these probes validated, we created adeno-associated viruses (AAVs) for BioID2-Synapsin and BioID2 as a negative control, and then injected them into the brains of newborn mice (Figure 1B). After weaning and supplying exogenous biotin via injections, biotinylated proteins were collected from purified cortical and hippocampal synaptosomes and analyzed using ultraperformance liquid chromatography-tandem mass spectrometry (UPLC-MS/MS) with label-free quantitation. Based on peptide identity, a total of 518 proteins were identified in all samples, which were then filtered based on fold enrichment over negative control and adjusted p-value (Figure 1C). This resulted in a network of 200 proteins selectively enriched in presynaptic terminals (Figure 1D).

Bioinformatic network analysis revealed that the Synapsin iBioID proteome is highly enriched for proteins implicated in presynaptic function (Figure 1E). Multiple compartments of presynaptic terminals were represented, including synaptic vesicles (20 proteins), active zones (8 proteins), and recycling endosomes (6 proteins). The proteome covered both excitatory and inhibitory terminals, as suggested by the identification of *Slc17a6* (Vglut2), *Slc1a2* (Glt1), *Slc32a1* (Vgat), and *Gad2.* DAVID analysis (Dennis et al., 2003) of the proteome found a significant enrichment for the biological processes of “synaptic vesicle endocytosis” (22 proteins, p=1.7×10^−6^) and “synaptic vesicle exocytosis” (30 proteins, p=3.6×10^−9^), among others. Eight proteins were of unknown function, not including the previously uncharacterized *Kiaa1107* (APache) which was recently shown to be involved in synaptic vesicle trafficking (Piccini et al., 2017). The only protein in the network strongly associated with the postsynaptic density (PSD) was Shank1, but there is recent evidence that Shank proteins have an unappreciated presynaptic function (Wu et al., 2017).

Regulators of the actin cytoskeleton were heavily overrepresented in the Synapsin iBioID proteome (54 proteins, p=9.8×10^−7^). Importantly, very few of these actin signaling molecules had been previously studied in presynaptic terminals (Figure 1E, Actin Cytoskeleton vs Known Presynaptic). The network also contained regulators of the microtubule and septin cytoskeleton, suggesting the capture of multiple components of the presynaptic cytomatrix. Overall, the network was highly interconnected with 54% of proteins (108 proteins) previously known to be presynaptic, suggesting high coverage of the presynaptic compartment.

To validate the Synapsin iBioID proteome, we selected 23 candidate genes that had not previously been shown to localize to presynaptic terminals, with a particular focus on actin regulators and proteins of unknown function (Table S1). We determined the localization of these proteins using Homology-Independent Universal Genome Engineering (HiUGE) (Gao et al., 2019), a CRISPR/Cas9-based technology to tag endogenous proteins. Hippocampal neurons were cultured from *H11Cas9* mice constitutively expressing Cas9 and then infected with AAVs for candidate C-terminal guide RNAs and their corresponding 2xHA-V5-Myc epitope-tag HiUGE donor (Figure 2A). Positive labeling was observed from 19 out of 23 genes, of which 14 displayed a robust signal with good signal-to-noise ratio above background fluorescence (Table S1).

**Figure 2.**
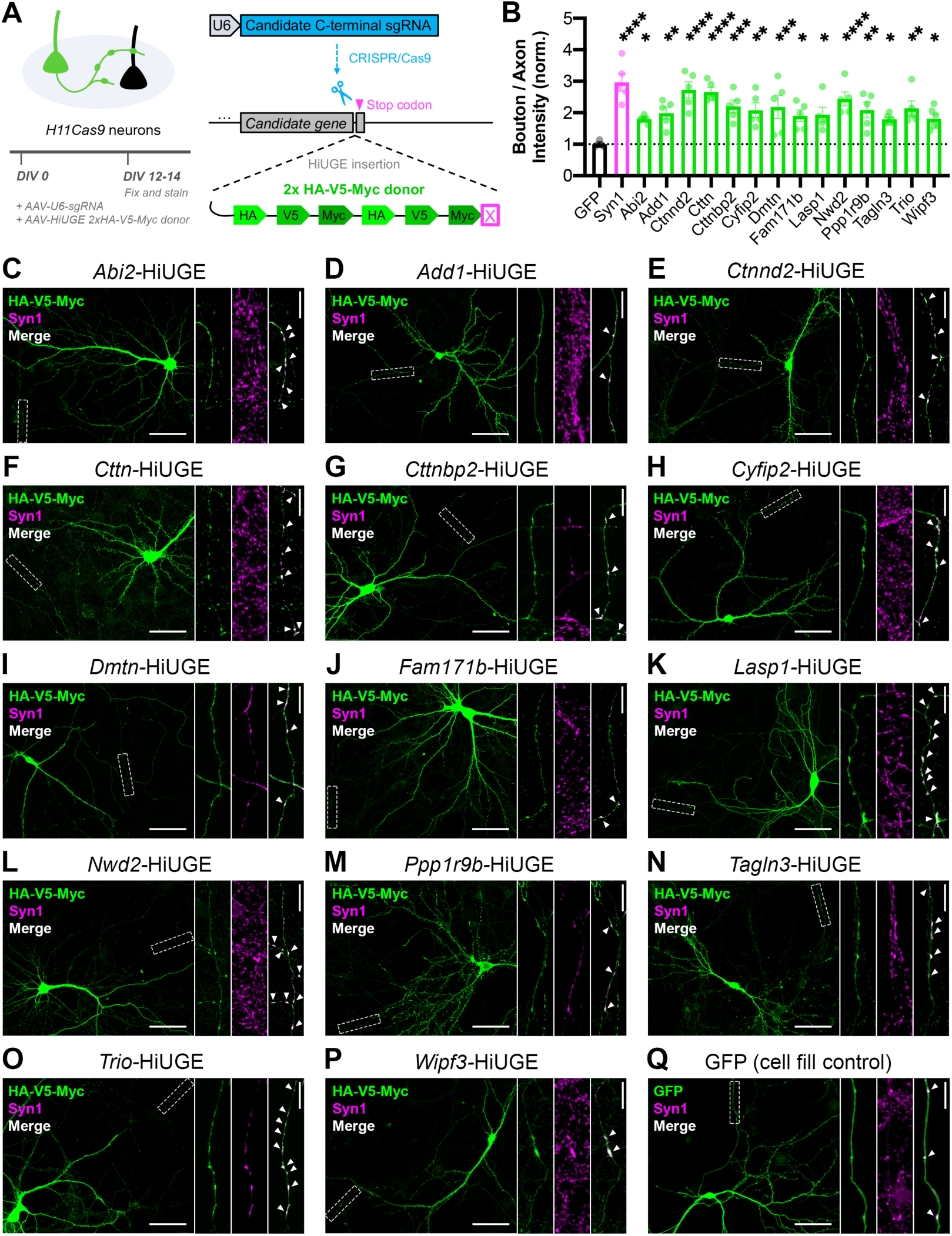
Validation of the presynaptic localization of Synapsin iBioID proteins. **(A)** Schematic of approach to tag endogenous proteins in neurons using HiUGE. Cultured hippocampal neurons were infected on DIV0 with AAVs containing the candidate sgRNA and a 2x-HA-V5-Myc HiUGE donor in the corresponding open reading frame. Neurons expressing a GFP cell fill were used as a control. **(B)** Quantification of presynaptic enrichment for GFP control (n=6 neurons), presynaptic marker Syn1 (Synapsin1, n=5), and candidate proteins (Abi2 n=6, Add1 n=5, Ctnnd2 n=5, Cttn n=5, Cttnbp2 n=5, Cyfip2 n=5, Dmtn n=5, Fam171b n=5, Lasp1 n=5, Nwd2 n=5, Ppp1r9b n=5, Tagln3 n=6, Trio n=5, Wipf3 n=6); one-way ANOVA (F_15,68_=5.401, p<0.0001) with Dunnett’s multiple comparisons test vs GFP: Syn1 (p<0.0001), Abi2 (p=0.0422), Add1 (p=0.0088), Ctnnd2 (p<0.0001), Cttn (p<0.0001), Cttnbp2 (p=0.0008), Cyfip2 (p=0.0032), Dmtn (p=0.0010), Fam171b (p=0.0215), Lasp1 (p=0.0156), Nwd2 (p<0.0001), Ppp1r9b (p=0.0030), Tagln3 (p=0.0437), Trio (p=0.0016), Wipf3 (p=0.0359). **(C-Q)** Representative images of the localization of candidate proteins (HA/V5/Myc or GFP; green) and a presynaptic marker (Synapsin1; magenta). Scale bars, 50μm. Insets show staining along axons. The merged image contains only Synapsin1 puncta within the axon, and white arrows point to presynaptic terminals (colocalized puncta). Scale bars, 5μm. All data are mean ± SEM. *p<0.05, **p<0.01, ***p<0.001, ****p<0.0001.

These 14 candidates included 12 actin regulators and 2 genes of unknown function, *Fam171b* and *Nwd2.* All endogenous candidate proteins were expressed throughout the cell body, dendrites, and in some cases dendritic spines (Figure 2C-P). As expected, all 14 proteins were also expressed in axons, with significant enrichment in presynaptic terminals as compared to a GFP cell fill (Figure 2B,Q). Together, this highlights the discovery of a considerable number of proteins that were previously not known to localize to presynaptic terminals, and suggests that the Synapsin iBioID network can reveal novel insights into presynaptic function.

### Diversity of presynaptic actin signaling and convergence on the Rac1-Arp2/3 pathway

On closer examination of the 54 actin cytoskeleton proteins in the Synapsin iBioID network, we uncovered a surprisingly rich diversity of actin signaling molecules in presynaptic terminals (Figure 3A). Many were adaptor proteins that linked the actin cytoskeleton to other signaling pathways or cellular structures, including endocytosis, phosphoinositide signaling, Arf GTPases, Rap GTPases, focal adhesions, and adherens junctions. At the level of actin monomers and filaments, we identified regulators involved in bundling and cross-linking filaments, severing filaments, capping filaments, and sequestering monomers. Of note, we found 2 proteins, *Tagln3* and *Wipf3,* known to bind actin but with uncharacterized cellular function.

**Figure 3.**
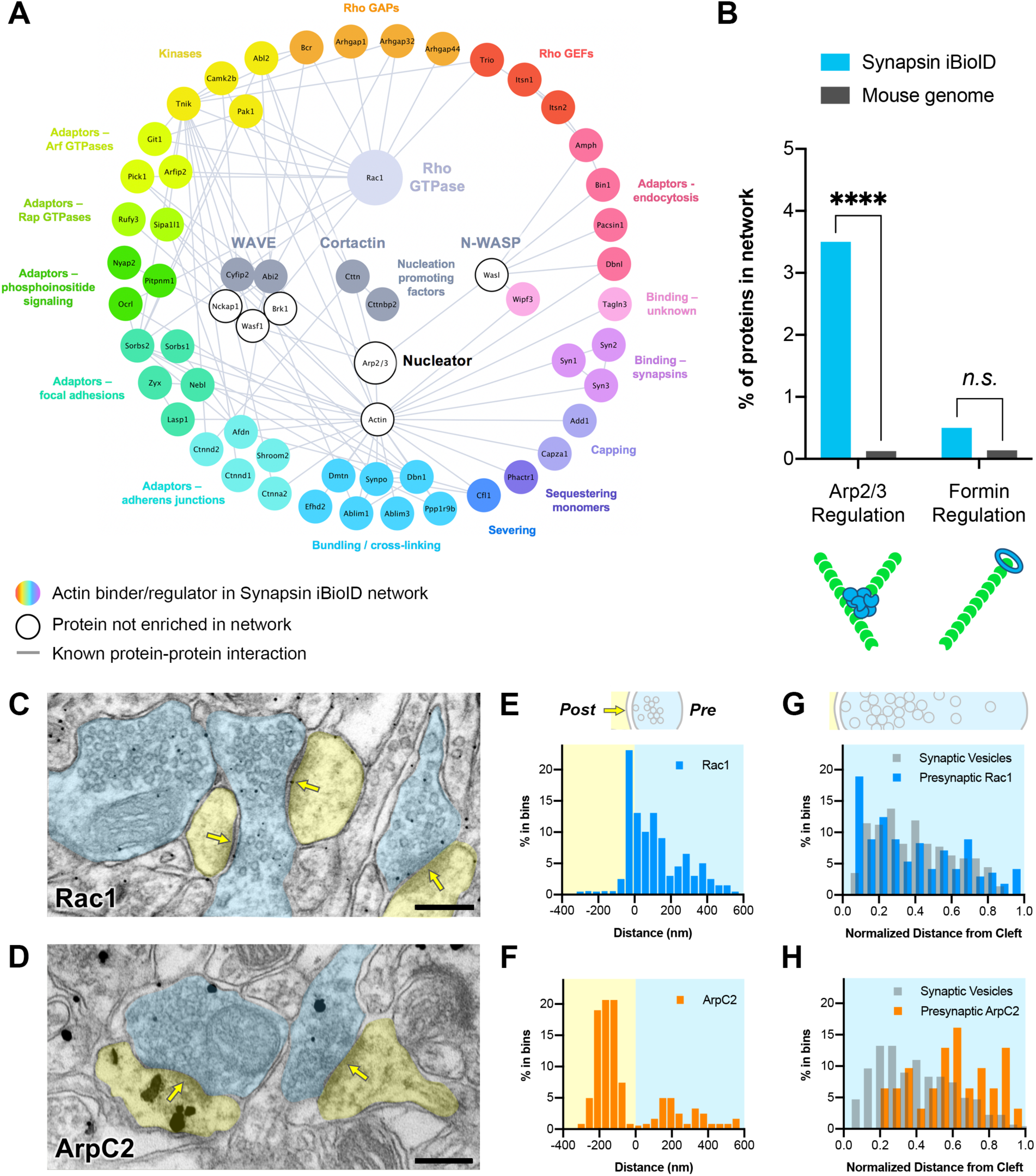
Actin signaling pathways in presynaptic terminals. **(A)** Network showing the diversity of presynaptic actin signaling pathways in the Synapsin iBioID proteome. Node titles correspond to gene name, and node size emphasizes the proteins further studied. Colored nodes are actin regulators in the Synapsin iBioID network, while white nodes are proteins not enriched compared to negative control. Edges are previously reported protein-protein interactions in the HitPredict database or by hand annotation. **(B)** Regulators of actin nucleation in the Synapsin iBioID network converge on Arp2/3, which nucleates branched actin filaments, rather than on formins, which nucleate linear actin filaments; FDR-adjusted hypergeometric test on Synapsin iBioID vs mouse genome for Arp2/3 regulation (p=1.2×10^−8^) and formin regulation (p=0.2555). **(C-D)** Representative pre-embedding immunogold-labeled electron micrographs in mouse hippocampal CA1 for **(C)** Rac1 and **(D)** ArpC2. Dendritic spines are pseudocolored yellow, presynaptic terminals are pseudocolored blue, and a yellow arrow points to the synaptic cleft. Scale bars, 200nm. **(E-F)** Axodendritic distribution of gold particles at the synapse coding for **(E)** Rac1 (blue, n=15 synapses) and **(F)** ArpC2 (orange, n=19 synapses). **(G-H)** Presynaptic distribution of synaptic vesicles (gray) and gold particles for **(G)** Rac1 (blue) and **(H)** ArpC2 (orange). Distances were normalized from the synaptic cleft based on the axodendritic length of the presynaptic terminal. ****p<0.0001, *n.s.* not significant.

Most interestingly, at the level of Rho GTPase signaling, only *Rac1* was significantly enriched. We also identified several Guanine Nucleotide Exchange Factors (GEFs: *Trio, Itsn1,* and *Itsn2)* and GTPase Activating Proteins (GAPs: *Bcr, Arhgap1, Arhgap32,* and *Arhgap44),* which activate and inactivate Rho GTPases, respectively. Downstream of Rac1, we identified its effector proteins *Pak1, Cttn,* and members of the WAVE complex *(Cyfip2* and *Abi2).* Cortactin and WAVE are nucleation promoting factors that activate the Arp2/3 complex to nucleate branched actin filaments. Using overrepresentation analysis, we found that regulators of Arp2/3, including Rac1, were significantly enriched in the Synapsin iBioID network (Figure 3B). In contrast, regulators of formins, which nucleate linear actin filaments (Schonichen and Geyer, 2010), were not significantly enriched. Thus, we hypothesized that Rac1-Arp2/3 signaling and branched actin play an important role in presynaptic terminals.

### Rac1 and Arp2/3 are associated with synaptic vesicle membranes *in vivo*

To validate the presence of Rac1 and Arp2/3 in presynaptic terminals *in vivo,* we investigated their localization using immunogold electron microscopy. We probed hippocampal CA1 of adult mice with antibodies against Rac1 and ArpC2, one of the non-actin-binding subunits of the Arp2/3 complex (Figure 3C-D). Rac1 localized to the PSD (Figure 3E), which is consistent with its known function in dendritic spine development and plasticity. However, unexpectedly, the majority of Rac1 labeling (70.3%) localized to presynaptic terminals and was closely associated with synaptic vesicle membranes. Gold particles coding for Rac1 were also located on plasma membranes. Overall, the distribution of presynaptic Rac1 was not significantly different from that of synaptic vesicles (Figure 3G).

As reported previously (Racz and Weinberg, 2008), ArpC2 was concentrated in dendritic spines approximately 200nm below the PSD (Figure 3F). However, a fraction of gold particles (25.6%) localized to presynaptic terminals with a consistent and specific distribution. ArpC2 was present among synaptic vesicles, although it preferentially localized to the presynaptic membrane beyond the synaptic vesicle cluster (Figure 3H). Very little immunolabeling was observed when the primary antibody was omitted as a negative control. In the few synapses that did have staining (<1%), there was diffuse non-specific signal across the synapse (Figure 3—figure supplement 1).

Taken together, the overlapping distributions of Rac1 and Arp2/3 at synaptic vesicles suggest a potential common presynaptic function related to synaptic vesicle modulation.

### Presynaptic Rac1 negatively regulates synaptic vesicle replenishment

We next tested whether Rac1 played a role in regulating neurotransmitter release. Since Rac1 functions postsynaptically in both development and plasticity (Hedrick and Yasuda, 2017; Tolias et al., 2011), we isolated its presynaptic function by using a mixed hippocampal culture system where presynaptic wildtype (WT) or knock-out (KO) neurons expressed channelrhodopsin (ChR2), and light-evoked responses were recorded from postsynaptic WT neurons (Figure 4A). To accomplish this, WT neurons were electroporated with tdTomato and then sparsely seeded with *Rac1^fl/fl^* neurons expressing ChR2-EYFP. To minimize developmental effects, AAV-hSyn-Cre was added after 10 days *in vitro* (DIV10) to half the coverslips, deleting *Rac1* from neurons expressing ChR2. In cultures without Cre, neurons expressing ChR2 remained functionally WT (Figure 4B).

**Figure 4.**
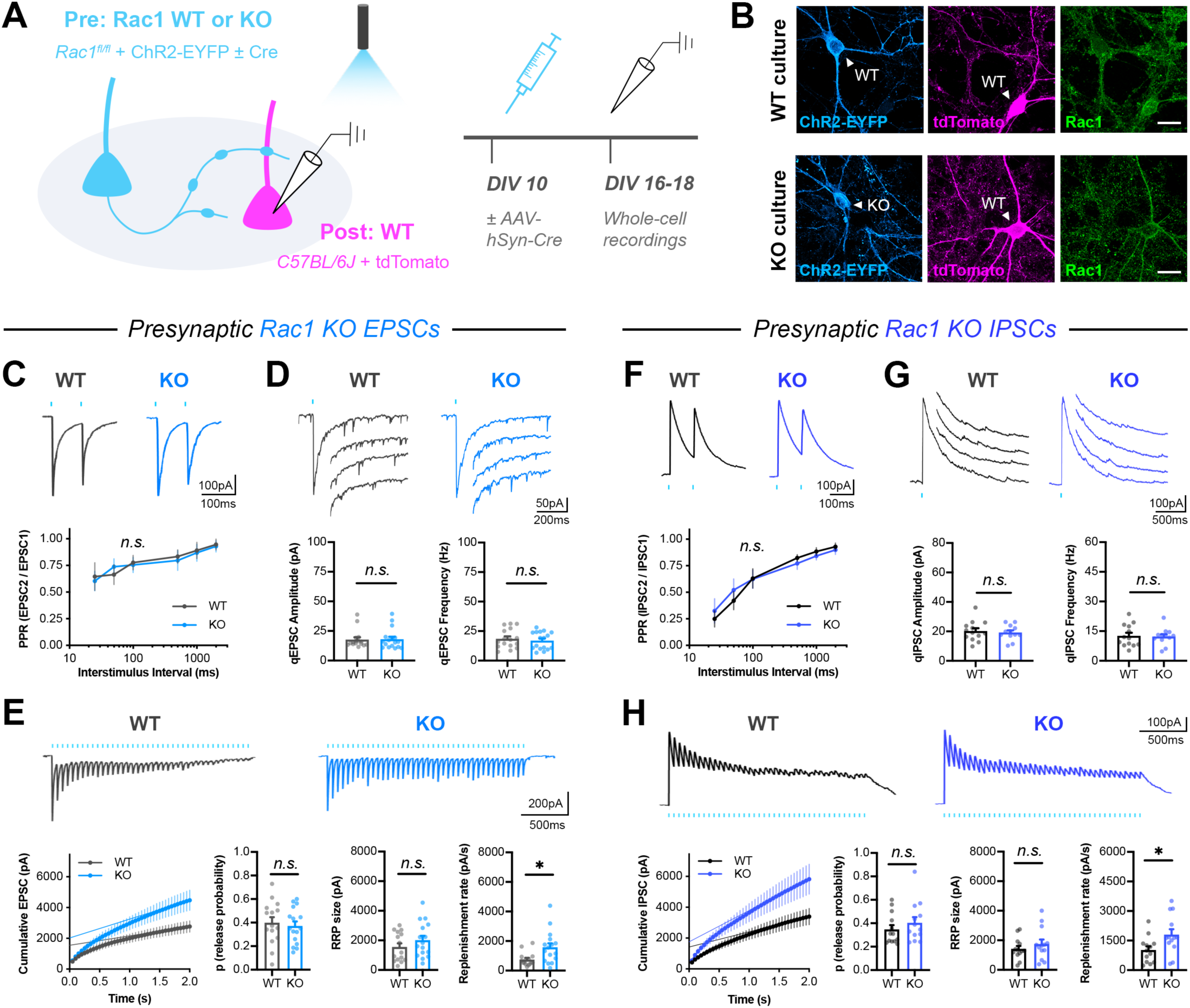
Presynaptic Rac1 negatively regulates synaptic vesicle replenishment. **(A)** Schematic of mixed hippocampal neuron cultures to isolate effects of presynaptic *Rac1* knockout. Whole-cell patch clamp recordings were conducted on tdTomato+ WT neurons with light delivered through the objective by a 460nm LED. (Representative images of WT and KO cultures fixed on DIV16 and stained for ChR2-EYFP (blue), tdTomato (magenta), and Rac1 (green). Scale bars, 15μm. **(C-E)** Light-evoked EPSCs in WT and KO cultures. Representative traces and quantification for: **(C)** PPR (WT n=15 neurons/3 cultures, KO n=17/3); two-way repeated measures ANOVA (F_1,30_=0.1462, p=0.7049) with Sidak’s multiple comparisons test: 25ms (p=0.9941), 50ms (p=0.7469), 100ms (p=0.9974), 500ms (p=0.9842), 1000ms (p=0.9989), 2000ms (p=0.9976). **(D)** Strontium-evoked qEPSCs (WT n=16/3, KO n=17/3); Mann-Whitney U tests for amplitude (U=130, p=0.8451) and frequency (U=120, p=0.5814). **(E)** 20Hz stimulation trains (WT n=15/3, KO n=16/3) and release probability (t-test, t_29_=0.4671, p=0.6439), RrP size (t-test, t_29_=1.271, p=0.2137), and replenishment rate (t-test, t_29_=2.574, p=0.0154). **(F-H)** Light-evoked IPSCs in WT and KO cultures. Representative traces and quantification for: **(F)** PPR (WT n=12/3, Ko n=11/3); two-way repeated measures ANOVA (F_1,21_=0.04765, p=0.8293) with Sidak’s multiple comparisons test: 25ms (p=0.8116), 50ms (p=0.5124), 100ms (p>0.9999), 500ms (p=0.4174), 1000ms (p=0.4110), 2000ms (p=0.7703). **(G)** Strontium-evoked qIPSCs (WT n=13/3, KO n=12/3); t-tests for amplitude (t_23_=0.2064, p=0.6798) and frequency (t_23_=0.2064, p=0.8383). (h) 20Hz stimulation trains (WT n=13/3, KO n=13/3) and release probability (t-test, t_24_=0.9657, p=0.3438), RRP size (t-test, t_24_=0.9253, p=0.3640), and replenishment rate (t-test, t_29_=2.382, p=0.0255). All data are mean ± SEM. *p<0.05, *n.s.* not significant.

We conducted whole-cell patch-clamp recordings from tdTomato-expressing WT neurons on DIV16-18, using light stimulation of presynaptic WT or KO neurons to evoke excitatory postsynaptic currents (EPSCs). Presynaptic *Rac1* deletion did not affect the amplitude, charge transfer, or kinetics of single evoked EPSCs (Figure 4—figure supplement 1C). It also did not affect the paired pulse ratio (PPR) (Figure 4C). Next, to assess quantal release parameters, EPSCs were evoked in the presence of Sr^2+^ (in place of Ca^2+^), which induces asynchronous quantal events after an initial synchronous release. Presynaptic *Rac1* deletion did not affect the amplitude or frequency of quantal events (Figure 4D). Finally, a 20Hz high frequency stimulation (HFS) train was used to probe synaptic vesicle recycling. Surprisingly, presynaptic *Rac1* deletion reduced short-term synaptic depression in response to HFS (Figure 4E). There was no effect on asynchronous release during the train, as measured by the steady-state basal current (Figure 4— figure supplement 1A,D). Quantification of the cumulative EPSC curve showed that presynaptic *Rac1* deletion increased the synaptic vesicle replenishment rate, without altering the release probability or size of the readily releasable pool (RRP).

To determine whether this was a common function of Rac1 across different kinds of presynaptic terminals, we next tested the effects of *Rac1* deletion from presynaptic inhibitory neurons. Inhibitory neurons were also present in our cultures, so we used light to evoke inhibitory postsynaptic currents (IPSCs). Presynaptic *Rac1* deletion from inhibitory neurons caused similar effects as in excitatory neurons. There were no effects on single evoked IPSCs (Figure 4—figure supplement 1E), PPR (Figure 4F), quantal events (Figure 4G), or asynchronous release during HFS trains (Figure 4—figure supplement 1 B,F). However, there was a reduction in the short-term depression of IPSCs due to an increase in the synaptic vesicle replenishment rate (Figure 4H). Together, these data suggest that Rac1 negatively regulates synaptic vesicle replenishment at both excitatory and inhibitory synapses.

### Presynaptic Arp2/3 negatively regulates release probability and vesicle replenishment

We next tested whether Arp2/3 has similar functions in regulating neurotransmitter release, since we found components of the WAVE complex in the presynaptic cytomatrix that are known to activate Arp2/3 downstream of Rac1. Using a similar mixed culture strategy, WT neurons were sparsely seeded with *ArpC3^fl/fl^;Ai14* neurons expressing ChR2-EYFP *(ArpC3* encodes a critical subunit of the Arp2/3 complex, and *Ai14* is a Cre reporter allele expressing tdTomato). Cre was added to half the coverslips on DIV10, and then whole-patch clamp recordings were conducted from non-fluorescent WT neurons on DIV16-18 (Figure 5A-B).

**Figure 5.**
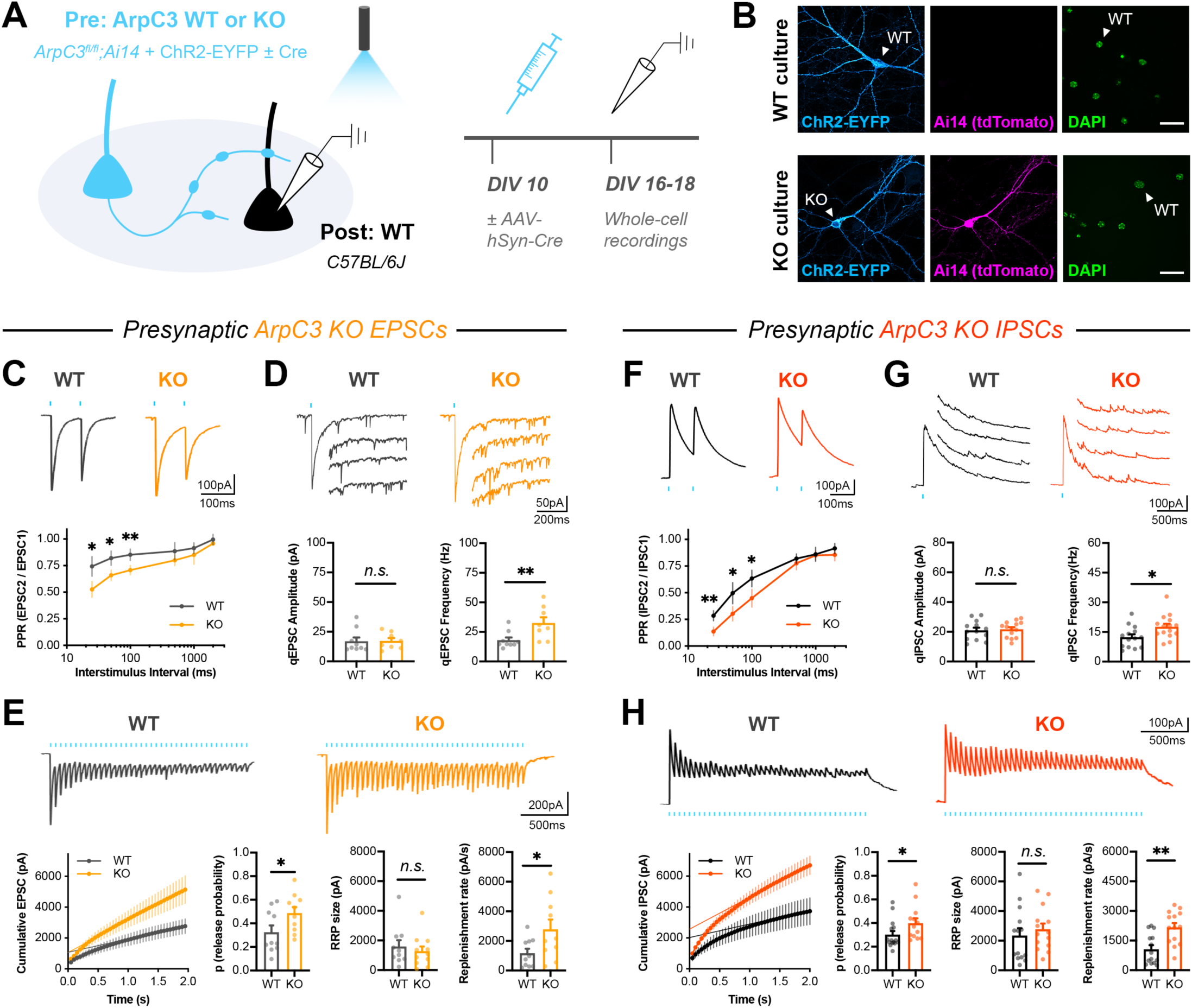
Presynaptic Arp2/3 negatively regulates release probability and synaptic vesicle replenishment. **(A)** Schematic of mixed hippocampal neuron cultures to isolate effects of presynaptic *ArpC3* knockout. Whole-cell patch clamp recordings were conducted on non-fluorescent WT neurons with light delivered through the objective by a 460nm LED. **(B)** Representative images of WT and KO cultures fixed on DIV16 and stained for ChR2-EYFP (blue), tdTomato (magenta), and DAPI (green). Scale bars, 25μm. **(C-E)** Light-evoked EPSCs in WT and KO cultures. Representative traces and quantification for: **(C)** PPR (WT n=10 neurons/3 cultures, KO n=12/3); two-way repeated measures ANOVA (F_1,20_=22.50, p=0.0001) with Sidak’s multiple comparisons test: 25ms (p=0.0435), 50ms (p=0.0194), 100ms (p=0.0099), 500ms (p=0.2168), 1000ms (p=0.2319), 2000ms (p=0.6130). **(D)** Strontium-evoked qEPSCs (WT n=9/3, KO n=8/3); Mann-Whitney U test for amplitude (U=31, p=0.6730) and t-test for frequency (t_15_=2.973, p=0.0095). **(E)** 20Hz stimulation trains (WT n=10/3, KO n=10/3) and release probability (t-test, t_18_=2.107, p=0.0494), RRP size (t-test, t_18_=0.3957, p=0.3957), and replenishment rate (t-test, t_18_=2.215, p=0.0399). **(F-H)** Light-evoked IPSCs in WT and KO cultures. Representative traces and quantification for: **(F)** PPR (WT n=14/3, KO n=13/3); two-way repeated measures ANOVA (F_1,25_=16.41, p=0.0004) with Sidak’s multiple comparisons test: 25ms (p=0.0022), 50ms (p=0.0117), 100ms (p=0.0111), 500ms (p=0.4100), 1000ms (p=0.9999), 2000ms (p=0.3992). **(G)** Strontium-evoked qIPSCs (WT n=14/3, KO n=15/3); t-tests for amplitude (t_27_=0.3989, p=0.6931) and frequency (t_27_=2.471, p=0.0201). **(H)** 20Hz stimulation trains (WT n= 14/3, KO n=14/3) and release probability (Mann-Whitney U-test, U=52, p=0.0350), RrP size (t-test, t_26_=0.6733, p=0.5067) and replenishment rate (t-test, t_26_=3.621, p=0.0012). All data are mean ± SeM. *p<0.05, **p<0.01, *n.s.* not significant.

Presynaptic *ArpC3* deletion in excitatory neurons increased the amplitude, charge, and decay time constants of single evoked EPSCs (Figure 5—figure supplement 1A). It also decreased PPR across interstimulus intervals (Figure 5C), suggesting an increased release probability. Presynaptic *ArpC3* deletion did not affect quantal amplitude, but it significantly increased the frequency of quantal events (Figure 5D). Since *ArpC3* deletion did not affect the density of synapses formed by axons (Figure 5—figure supplement 2), the frequency effect was likely due to increased release probability rather than increased synapse number. Presynaptic *ArpC3* deletion also reduced short-term synaptic depression in response to 20Hz light stimulation (Figure 5E), with no significant change in asynchronous release during the train (Figure 5—figure supplement 1B). Quantification of the cumulative EPSC showed that there was an increase in both release probability and synaptic vesicle replenishment rate. The same phenotypes were observed by *ArpC3* deletion in presynaptic inhibitory neurons (Figure 5F-H, Figure 5—figure supplement 1C-D).

Importantly for these experiments, both *ArpC3* and *Rac1* WT and KO neurons were able to consistently fire light-evoked action potentials at 20Hz (Figure 5—figure supplement 3A-B, F-G). *ArpC3* deletion did not affect most intrinsic properties of neurons, but it did increase the width of light-evoked action potentials (Figure 5—figure supplement 3H-I). It also increased the width of action potentials from current injection (Figure 5—figure supplement 3J), suggesting there was a change in intrinsic membrane properties. Because of this, it is possible that the effect of *ArpC3* deletion on synaptic vesicle replenishment, as seen through increased current amplitudes at the end of the 20Hz train, was actually caused by an increased action potential width or increased release probability during each stimulation. However, since *Rac1* deletion increased synaptic vesicle replenishment rate without affecting the action potential waveform (Figure 5—figure supplement 3C-E) or release probability, this strongly suggests that these phenotypes are separable, and that the Rac1-Arp2/3 pathway functions to negatively regulate synaptic vesicle replenishment at both excitatory and inhibitory synapses.

### Bidirectional control of presynaptic Rac1 signaling modulates short-term depression

We next set out to test whether acute modulation of Rac1 signaling would similarly affect synaptic vesicle replenishment. To accomplish this, we utilized photoactivatable Rac1 (PA-Rac1) constructs with dominant negative (DN) or constitutively active (CA) Rac1 mutations (Wu et al., 2009), along with additional mutations in the photoactivation domain to decrease background activity in the dark (Hayashi-Takagi et al., 2015). PA-Rac1 constructs were co-expressed with the red-shifted opsin ChrimsonR (Klapoetke et al., 2014) by fusing them with a P2A ribosome skip sequence along with an HA epitope tag (Figure 6A). This allowed for dual-color, light-driven control of both Rac1 signaling and neurotransmitter release in the same presynaptic neurons. Cultured hippocampal neurons were sparsely seeded with neurons expressing the ChrimsonR-tdTomato-P2A-HA-PA-Rac1 DN or CA constructs, or ChrimsonR-tdTomato alone as the WT control. Both ChrimsonR and PA-Rac1 expressed readily in the same neurons.

**Figure 6.**
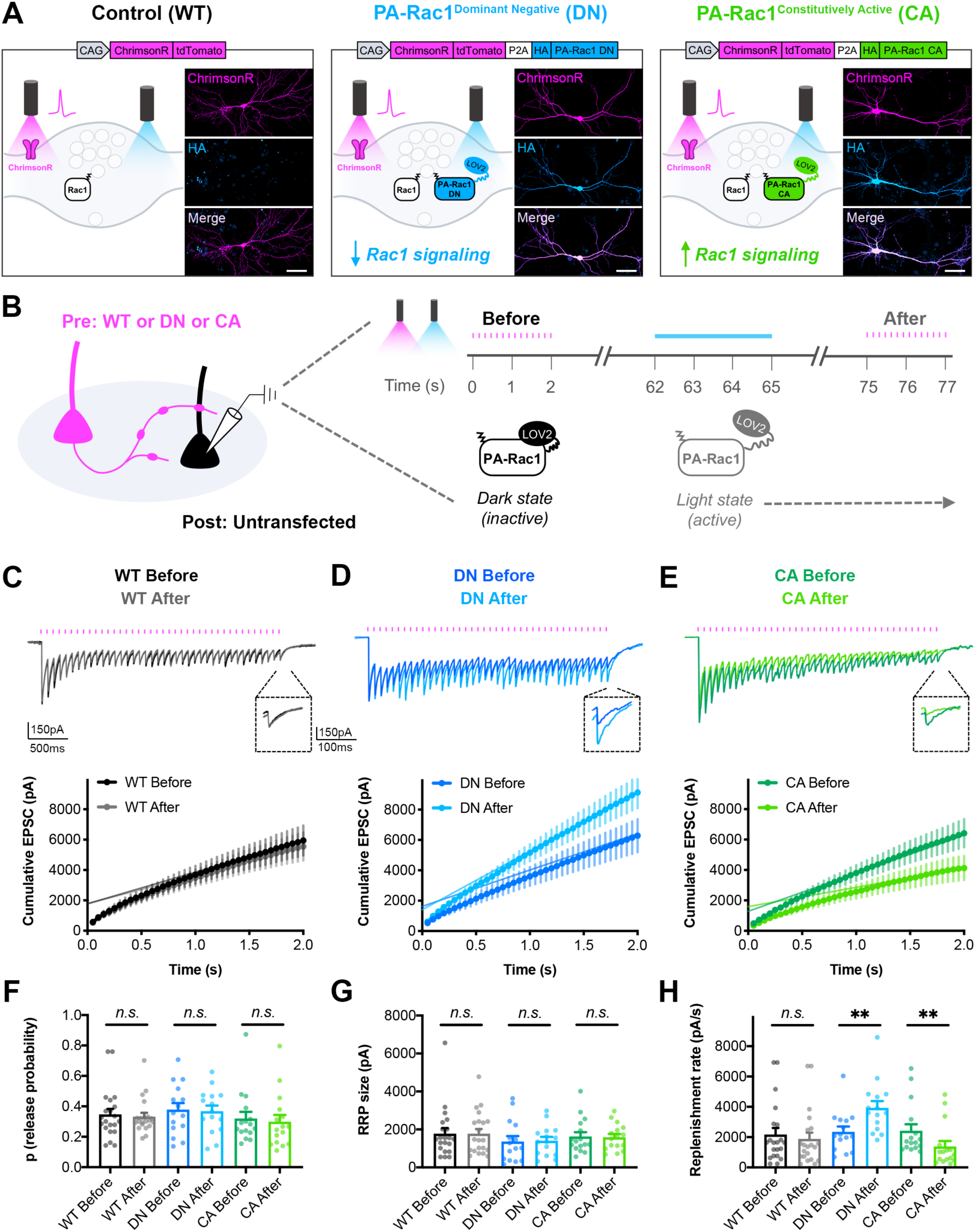
Bidirectional control of presynaptic Rac1 signaling modulates short-term synaptic depression. **(A)** Schematic of constructs created to control the firing of presynaptic neurons with reduced or enhanced Rac1 signaling. ChrimsonR-tdTomato was expressed alone as a control (WT), or co-expressed with HA-tagged photoactivatable Rac1 (PA-Rac1) with dominant negative (DN) or constitutively active (CA) mutations. Insets are representative images of WT, DN, and CA cultures fixed on DIV14 and stained for tdTomato (magenta) and HA (blue). Scale bars, 50μm. **(B)** Schematic of experimental design. Whole-cell patch clamp recordings were conducted on non-fluorescent neurons with light delivered through the objective by an LED. The “Before” 20Hz train was evoked by 525-660nm light. After waiting one minute for recovery, PA-Rac1 was brought into the open configuration by 460nm light to modulate presynaptic Rac1 signaling. Then, the “After” 20Hz train was evoked by 525-660nm light. **(C-E)** Representative traces and quantification of before and after EPSC trains in **(C)** WT cultures (black, gray, n=21 neurons/3 cultures), **(D)** DN cultures (blue, cyan, n=15/3), and **(E)** CA cultures (green, lime, n=16/3). **(F-H)** Estimates from cumulative EPSCs in all cultures of: **(F)** Release probability; WT (Mann-Whitney U-test, U=217, p=0.9355), DN (t-test, t_28_=0.1803, p=0.8582), CA (Mann-Whitney U-test, U=108, p=0.4677). **(G)** RRP size; WT (Mann-Whitney U-test, U=217, p=0.9355), DN (t-test, t_28_=0.1081, p=0.9147), and CA (Mann-Whitney U-test, U=124, p=0.8965). **(H)** Replenishment rate; WT (Mann-Whitney U-test, U=182, p=0.3394), DN (t-test, t_28_=2.800, p=0.0092), cA (Mann-Whitney U-test, U=48, p=0.0019). All data are mean ± SEM. **p<0.01, *n.s.* not significant.

On DIV14-16, whole-cell patch-clamp recordings were conducted from non-fluorescent postsynaptic neurons in the dark (Figure 6B). A 20Hz train was evoked with red-shifted light to obtain the baseline EPSC response. After waiting one minute for recovery, blue light was used to stimulate PA-Rac1 into the light state, where it remained on the order of seconds to minutes before decaying back to the dark, closed state (Wang et al., 2016). In the light state, PA-Rac1 was able to act in a DN or CA manner to modulate Rac1 signaling, and a second 20Hz train was quickly evoked with red-shifted light to determine the effect. ChrimsonR, although red-shifted, is known to still be activated by blue light, so light intensities were chosen to minimize crosstalk. Any remaining crosstalk did not have an effect on WT control neurons, since the EPSC trains before and after blue light stimulation were not significantly different (Figure 6C).

Acute inactivation of presynaptic Rac1 signaling phenocopied the genetic deletion; presynaptic stimulation of PA-Rac1 DN resulted in reduced short-term synaptic depression due to an increase in synaptic vesicle replenishment rate (Figure 6D, F-H). Conversely, acute activation of Rac1 signaling drove the phenotype in the opposite direction. Presynaptic stimulation of PA-Rac1 CA resulted in increased short-term synaptic depression due to a decrease in synaptic vesicle replenishment rate (Figure 6E, F-H). Neither manipulation affected release probability or RRP size. This bidirectional effect demonstrates that presynaptic Rac1 signaling sets the precise level of synaptic depression through its negative regulation of vesicle replenishment.

### Action potential trains activate Rac1 in presynaptic terminals

To investigate the dynamics of presynaptic Rac1 signaling, and to determine whether its activity is coupled to action potential trains, we used 2-photon Fluorescence Lifetime Imaging Microscopy (2pFLIM) in conjunction with a FRET-based sensor of Rac1 activity (Hedrick et al., 2016; Takahashi et al., 2015). AAVs encoding the FLIM donor (mEGFP-Rac1) and FLIM acceptor (mCherry-Pak2 GTPase Binding Domain-mCherry) were microinjected into CA3 of organotypic hippocampal slices on DIV10-13 (Figure 7A-B). After allowing at least 7 days for axonal expression, 2pFLIM was conducted on presynaptic boutons in CA1. A stimulation electrode was placed in the Schaffer collaterals at the CA3/CA1 border, and a recording electrode was placed in CA1 to record evoked field potentials.

**Figure 7.**
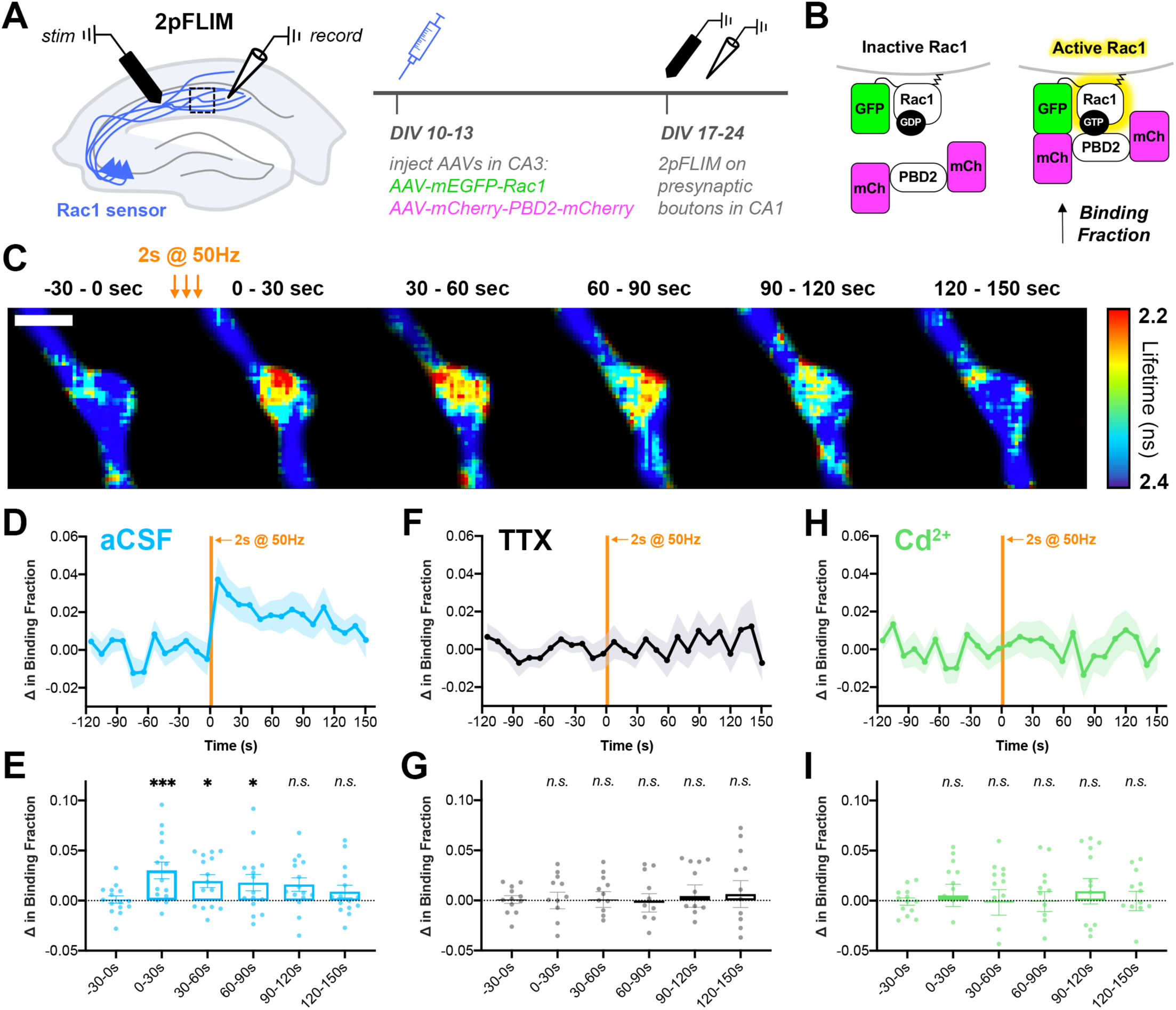
Action potential trains activate Rac1 in presynaptic terminals. **(A)** Experimental design in organotypic hippocampal slices. **(B)** Schematic of Rac1 sensor. Activation of Rac1 leads to its association with the GTPase binding domain of Pak2_^R71C,S78A^_ (PBD2), increasing FRET between GFP and mCherry. This is measured as a decrease in fluorescence lifetime, or an increase in binding fraction. **(C)** Representative 2pFLIM images of a bouton before and after stimulation for 2s at 50Hz. Scale bar, 1μΐΓ **(D)** Mean time course of the change in binding fraction of the Rac1 sensor in aCSF (cyan, n=15 boutons/5 slices) with **(E)** quantification; one-way repeated measures ANOVA (F_6,84_=3.89, p=0.0018) with Dunnett’s multiple comparisons test vs the baseline (−30–0s): 0-30s (p=0.0005), 30-60s (p=0.0102), 60-90s (p=0.0142), 90-120s (p=0.2881), 120-150s (p=0.6807). **(F)** Mean time course of Rac1 sensor in TTX (black, n=12/4) with **(G)** quantification; one-way repeated measures ANOVA (F_6,66_=0.8539, p=0.5334) with Dunnett’s multiple comparisons test vs the baseline (−30-0s): 0-30s (p=0.9930), 30-60s (p=0.9839), 60-90s (p=0.6430), 90-120s (p=0.7654), 120-150s (p=0.6548). **(H)** Mean time course of Rac1 sensor in Cd_^2^+_ (green, n=13/4) with **(I)** quantification; one-way repeated measures ANOVA (F_6,72_=0.2728, p=0.9479) with Dunnett’s multiple comparisons test vs the baseline (−30-0s): 0-30s (p>0.9999), 30-60s (p=0.9996), 60-90s (p=0.9996), 90-120s (p=0.9997), 120-150s (p=0.8896). All data are mean ± SeM. *p<0.05, ***p<0.001, *n.s.* not significant.

Upon electrical stimulation to induce action potential trains, Rac1 activity was significantly elevated in presynaptic boutons (Figure 7C-E). Interestingly, this increase in activity was persistent for a period of 60-90s, as measured by an increase in binding fraction. There was no change in binding fraction in the presence of TTX (Figure 7F-G), confirming that presynaptic Rac1 activation is action potential-dependent. There was also no change in binding fraction in the presence of Cd^2+^ (Figure 7H-I), demonstrating that presynaptic Rac1 activation requires calcium influx through voltage-gated calcium channels. Post-hoc staining of slices revealed that nearly all (∼92%) of boutons from GFP+ mCherry+ axons contained synapsin (identified by local axonal swelling, Figure 7—figure supplement 1), showing that these boutons tightly corresponded to presynaptic terminals. In summary, these data demonstrate a high-frequency train of action potentials leads to the activation of Rac1 in presynaptic terminals through calcium signaling. The time scale of Rac1 activity observed, on the order of tens of seconds, further supports its physiological role in presynaptic plasticity.

## DISCUSSION

Here, we used iBioID with a Synapsin probe to identify 200 proteins in cortical and hippocampal presynaptic terminals *in vivo,* with significant enrichment of cytoskeletal-associated proteins. This extends previous efforts to identify the proteome of isolated synaptic vesicles and active zone fractions (Abul-Husn et al., 2009; Boyken et al., 2013; Burre et al., 2006; Coughenour et al., 2004; Morciano et al., 2009; Morciano et al., 2005; Takamori et al., 2006; Weingarten et al., 2014; Wilhelm et al., 2014). Synapsin is thought to reside in multiple presynaptic terminal compartments (Guarnieri et al., 2015; Hilfiker et al., 1999), so the spread of activated biotin allowed for identification of proteins throughout these regions. Thus, while our iBioID approach identified components of the synaptic vesicles and the active zone, it also allowed for a more holistic view of presynaptic terminal space, including the presynaptic cytomatrix. Indeed, our analysis revealed a large number of proteins (92/200) that were not previously known to localize to presynapses, and these were mainly involved in actin cytoskeleton regulation, cell-cell adhesion, or other signaling pathways. We also validated the presynaptic localization of 14 of these proteins using an endogenous genomic tagging approach and an additional protein, Rac1, using electron microscopy. These results provide a new framework from which to view the cellular biology of presynaptic physiology and uncover a new actin-based mechanism of short-term plasticity. We anticipate that the technological advances we report here, combining electrophysiology with paired optogenetic control of both activity and signaling, will enable new insights into how these proteins function to couple neuronal spiking with transmitter release.

### Actin remodeling as a new mechanism of short-term synaptic depression

Although there is evidence suggesting the existence of active mechanisms to cause short-term depression, the identity of these processes remains unresolved (Bellingham and Walmsley, 1999; Byrne, 1982; Chen et al., 2004; Hsu et al., 1996; Kraushaar and Jonas, 2000; Parker, 1995; Sullivan, 2007; Thomson and Bannister, 1999; Waldeck et al., 2000; Xu and Wu, 2005; Zucker and Bruner, 1977). Our experiments using PA-Rac1 reveal that elevating or dampening levels of presynaptic Rac1 activity inversely alters synaptic vesicle replenishment rates, bidirectionally modulating the degree of short-term synaptic depression. Thus, regulation of Rac1 activity, upstream of Arp2/3-dependent actin polymerization, appears to play a central role in connecting activity to the tuning of vesicle replenishment, driving short-term depression. This pathway acts similarly at both excitatory and inhibitory synapses, suggesting it is a fundamental aspect of presynaptic function.

The mechanism by which this occurs likely does not depend on actin-synapsin interactions since synapsin function differs across cell types (Gitler et al., 2004a; Patzke et al., 2019). Signaling from Rac1 to Arp2/3 is known to nucleate branched actin filaments, which in presynaptic terminals may act as a barrier to diffusion to restrict synaptic vesicle mobility (Rothman et al., 2016). This would also make the active zone proteins Bassoon and Piccolo less available for accelerating vesicle replenishment (Butola et al., 2017; Hallermann et al., 2010). Alternatively, the Rac1-Arp2/3 pathway could negatively regulate synaptic vesicle endocytosis, although this would be surprising since Arp2/3 is required for endocytosis in yeast (Moreau et al., 1997) and actin itself is required for most, if not all forms of synaptic vesicle endocytosis in mammals (Soykan et al., 2017; Watanabe et al., 2013; Wu et al., 2016). This could also potentially explain why previous studies using actin depolymerizing agents did not detect an increase in synaptic vesicle replenishment. These pharmacological agents would have impaired the actin required for endocytosis (and thus synaptic vesicle replenishment), thereby masking forms of negative regulation by other pools of actin such as those we report here.

### Insights into the structure and function of the presynaptic actin cytoskeleton

Our systematic genetic analyses of Rac1 and Arp2/3 function provide new insights into the regulation of the presynaptic actin cytoskeleton that could not be understood using pharmacological approaches. Previously, it was thought that actin was not present within the synaptic vesicle cluster but rather localized around its periphery and at endocytic zones, based on studies using immunoelectron microscopy or cryoelectron tomography (Fernandez-Busnadiego et al., 2010; Pechstein and Shupliakov, 2010; Siksou et al., 2007). However, our finding that Rac1 and Arp2/3 are closely associated with vesicle membranes within the synaptic vesicle cluster suggests this may need to be re-examined. Since this pathway appears to be coupled to activity during short-term plasticity, we speculate that the actin filaments produced are too transient to be detected by conventional methods.

We found that Arp2/3-dependent actin plays a role not only in synaptic vesicle replenishment, but also in the negative regulation of release probability. Loss of Arp2/3 also led to a change in intrinsic membrane properties, because action potential width was increased by both ChR2 stimulation and current injection. Recently it was shown that Arp2/3-dependent actin slows the inactivation rate of Kv3.3, a voltage-gated potassium channel that is important for action potential repolarization (Zhang et al., 2016). Thus, it is plausible that loss of Arp2/3 could increase action potential width via impaired repolarization. Increased width of the action potential would also likely lead to an increase in calcium influx during repetitive stimulation, explaining the increase in release probability we observed.

Nonetheless, our results highlight that there are at least two different pools of branched actin in presynaptic terminals: a Rac1-Arp2/3-dependent pool in the synaptic vesicle cluster that regulates vesicle replenishment and synaptic depression, and an Arp2/3-dependent pool that regulates release probability independently of Rac1. As discussed earlier, there may also be a third pool of actin involved in synaptic vesicle endocytosis that is not dependent on Rac1 or Arp2/3. Multiple pools of actin assemblies existing in subdomains of presynaptic terminals is consistent with the diversity of actin regulators identified within the Synapsin iBioID proteome. Based on the identity of these proteins, it is now possible to use similar genetic analyses to delineate the presynaptic functions of actin severing proteins, bundling proteins, crosslinking proteins, and others during both baseline synaptic transmission and synaptic plasticity. It is particularly intriguing that presynaptic boutons enlarge after long-term potentiation in a form of structural plasticity (Chereau et al., 2017). We propose this new form of structural plasticity will be informed by the highly diverse nature of actin regulatory proteins enriched in presynaptic terminals, like that of the postsynapse. In support of this idea, actin remodeling was recently shown to be involved in a form of long-term depression at GABAergic terminals that is mediated by retrograde cannabinoid signaling (Monday et al., 2020).

### Rac1 signaling in presynaptic terminals and implications for neurological diseases

Postsynaptic Rho GTPase signaling is clearly important for dendritic spine development, maintenance, and plasticity (Hedrick and Yasuda, 2017; Tolias et al., 2011), but here we show that Rac1 is also abundant in presynaptic terminals, where it is involved in the essential processes of synaptic vesicle replenishment and short-term synaptic plasticity. We found that presynaptic Rac1 is transiently activated by calcium influx during HFS, but what is the upstream calcium sensor? It is tempting to speculate the involvement of CaMKII, since CaMKII is present in presynaptic terminals (Ding et al., 2013) and interacts with L-type voltage-gated calcium channels (Abiria and Colbran, 2010), and we detected CaMKIIp in the Synapsin iBioID proteome. Interestingly, the Rac1 GEF identified in our proteomics, Trio, is a likely CaMKII substrate important for plasticity at postsynaptic sites (Herring and Nicoll, 2016), and thus may also modulate Rac1 presynaptically. The conservation of Rac1 plasticity signaling at both the pre- and postsynapse is a surprising finding given the different mechanisms tuning efficacy between these sites. This highlights the concept that synaptic actin remodeling is a convergent mechanism for coupling activity to changes in the efficacy of neurotransmission regardless of synaptic locale.

Defects in Rho GTPases signaling pathways are also heavily implicated in neurodevelopmental disorders (Spence and Soderling, 2015; Yan et al., 2016), including missense mutations in *Rac1* that cause intellectual disability (Lelieveld et al., 2016; Reijnders et al., 2017) and an Arp2/3 mutation associated with schizophrenia (Gulsuner et al., 2020). Studies investigating the neural basis for these cognitive impairments, including our own, have focused mainly on deficits in dendritic spine development and plasticity with only limited assessments of presynaptic function (Kim et al., 2013; Kim et al., 2015; Soderling et al., 2007; Tian et al., 2018; Volk et al., 2015; Zoghbi and Bear, 2012). Our results compel a re-evaluation to include the potential presynaptic phenotypes in these diseases.

Together, this study sheds light on the previously uncharacterized and conserved regulation of presynaptic actin, and creates a new framework for understanding how presynaptic structure and strength may be altered during learning and disease. The Rac1-Arp2/3 pathway is a common regulator of plasticity at both sides of the synapse, and many other signaling pathways that are thought to be confined to postsynaptic sites may also be engaged presynaptically. The experimental strategies and resources that we developed here open numerous avenues of future research, and bring into focus the exquisite, complex signaling that occurs in presynaptic terminals.

## ACKNOWLEDGEMENTS

We thank George Augustine for generously providing the GFP-Synapsin1a construct. We also thank members of the Yasuda lab for 2pFLIM support, in particular David Kloetzer for coordinating visits to Florida, Jaime Richards for preparing organotypic slices, and Paula Parra-Bueno for extensive help setting up imaging experiments. We also thank members of the Soderling lab for helpful discussion and critical reviews of the manuscript. This work was supported by an NIH grant (R01MH103374) to S.H.S.; a European Union and European Social Fund grant (EFOP-3.6.2-16-2017-0008), an NKFIH grant (KKP126998), and a János Bolyai Research Fellowship of the Hungarian Academy of Sciences to B.R.; and an NSF Graduate Research Fellowship (DGE-1644868) to S.D.

## AUTHOR CONTRIBUTIONS

Conceptualization, S.D. and S.H.S.; Methodology, S.D., B.R., Y.G., E.J.S., R.Y., and S.H.S.; Investigation, S.D., B.R., W.E.B., and E.J.S.; Resources, B.R., E.J.S., R.Y., and S.H.S.; Writing - Original Draft, S.D. and S.H.S.; Writing - Review and Editing, S.D., B.R., W.E.B., E.J.S., Y.G., R.Y., and S.H.S.; Visualization, S.D.; Supervision, R.Y. and S.H.S.; Funding Acquisition, S.D., B. R., and S.H.S.

## COMPETING INTERESTS

S.H.S. and Y.G. have filed a patent application related to the HiUGE technology, and the IP has been licensed to CasTag Biosciences. S.H.S. is a founder of CasTag Biosciences. R.Y. is a founder and shareholder of Florida Lifetime Imaging LLC, a company that helps people set up FLIM.

## MATERIALS AND METHODS

### Animals

*C57BL/6J* mice (stock #000664) and *H11Cas9* mice (stock #028239) were purchased from The Jackson Laboratory. *Rac1^fl/fl^and ArpC3^fl/fl^;Ai14* mice have been described previously (Chrostek et al., 2006; Kim et al., 2015). Mice of both sexes were used for all experiments. All mice were housed (two to five mice per cage) in facilities provided by Duke University’s Division of Laboratory Animal Resources or Max Planck Florida Institute for Neuroscience’s Animal Resource Center. All experimental procedures were conducted with protocols approved by the Institutional Animal Care and Use Committee at Duke University and Max Planck Florida Institute for Neuroscience, in accordance with National Institutes of Health guidelines.

### Primary neuronal culture

Primary hippocampal neuron cultures were prepared from mice by isolating hippocampi from P0-P1 pups of both sexes under a dissection microscope. For mixed cultures, hippocampi were stored in Hibernate-A medium (Gibco) supplemented with 2% B-27 (Gibco) for 1-2 days at 4°C until the second litter was born. Then, hippocampi were incubated with papain (Worthington) at 37°C for 18 min, dissociated by gentle trituration, and plated onto 18mm glass coverslips treated with poly-L-lysine (Sigma). Electroporations were performed immediately before plating neurons using a Nucleofector 2b Device (Lonza) and the Mouse Neuron Nucleofector Kit (Lonza), following the manufacturer’s instructions. Neurons were maintained in Neurobasal A medium supplemented with 2% B-27 and 1% GlutaMAX (Gibco) in an incubator at 37°C and 5% CO_2_. After 5 days in culture, 5μΜ cytosine arabinoside (Sigma) was added to inhibit glial division. Subsequently, medium was half exchanged every 3-4 days. For PA-Rac1 experiments, cultures were wrapped in foil to minimize background activity due to ambient light.

### Organotypic hippocampal slice culture

Organotypic hippocampal slices were prepared from *C57BL/6J* mice. Briefly, P3-P8 pups of both sexes were euthanized by deep anesthesia with isoflurane followed by decapitation. Hippocampi were dissected from the brain, cut into coronal slices (350μm thickness) using a McIlwain tissue chopper (Ted Pella), and plated on Millicell hydrophilic PTFE membranes (Millipore). Slices were maintained in culture medium containing MEM medium (Life Technologies), 20% horse serum, 1mM L-glutamine, 1mM CaCh, 2mM MgSO_4_, 12.9mM D-glucose, 5.2mM NaHCO_3_, 30mM HEPES, 0.075% ascorbic acid, μg/ml insulin, and 1% penicillin-streptomycin. Medium was fully exchanged every 2-3 days.

### Plasmids

pCMV-EGFP-Synapsin1a (rat) was generously provided by George Augustine. pAAV-hSyn-hChR2 (H134R)-EYFP (Addgene plasmid #26973) was a gift from Karl Deisseroth. pCAG-ChrimsonR-tdTomato (Addgene plasmid #59169) was a gift from Edward Boyden. pCMV-mEGFP-Rac1 (Addgene plasmid #83950) and pCAG-mCherry-PBD2-mCherry (Addgene plasmid #83951) were a gift from Ryohei Yasuda. pAAV-hSyn-BioID2-HA, pAAV-hSyn-BioID2-Linker-BioID2-HA, pCAG-GFP, pAAV-hSyn-Cre, and pBetaActin-tdTomato were previously generated in the Soderling lab.

pAAV-hSyn-BioID2-Linker-Synapsin1a-HA was generated by PCR of Synapsin1a from pCMV-EGFP-Synapsin1a (primers FWD: 5’GGTGTCTAAGGAATTCAACTACCTGCGGCGC CGC3’ and REV: 5’AAGGGTAAGCGCTAGCGTCGGAGAAGAGGCTGGC3’) and insertion into the EcoRI/NheI sites of pAAV-hSyn-BioID2-Linker-BioID2-HA using In-Fusion cloning (TaKaRa). pCAG-ChrimsonR-tdTomato-P2A-HA-PA Rac1 (DN) and pCAG-ChrimsonR-tdTomato-P2A-HA-PA Rac1 (CA) were generated by synthesis of BioXp tiles (SGI-DNA) containing the C-terminus of tdTomato fused to P2A-HA-PA Rac1. PA Rac1 sequences (Wu et al., 2009) contained L514K and L531E mutations in the PA domain to reduce background activity in the dark (Hayashi-Takagi et al., 2015) as well as DN (T17N) or CA (Q61L, E91H, and N92H) mutations in Rac1. DNA tiles were inserted into the NotI/BsmBI sites of pCAG-ChrimsonR-tdTomato using In-Fusion cloning. pAAV-hSyn-mEGFP-Rac1 was generated by PCR of mEGFP-Rac1 from pCMV-mEGFP-Rac1 (primers FWD: 5’ACCGGC TAGAGTCGACACCATGGTGAGCAAGGG3’ and REV: 5’TAAGCGAATTGGATCCTTACA ACAGCAGG3’) and insertion into the SalI/BamHI sites of pAAV-hSyn-BioID2-Linker-BioID2-HA using In-Fusion cloning. pAAV-hSyn-mCherry-PBD2-mCherry was generated by PCR of mCherry-PBD2-mCherry from pCAG-mCherry-PBD2-mCherry (primers FWD: 5’ACCGGCTA GAGTCGACGGTCGCCACCATGGTGA3’ and REV: 5’TAAGCGAATTGGATCCGCGGCCG CTTACTTGTA3’) and insertion into the SalI/BamHI sites of pAAV-hSyn-BioID2-Linker-BioID2-HA using In-Fusion cloning. For generation of HiUGE plasmids, see the “HiUGE tagging of candidate genes” section below. All constructs generated in the Soderling lab were validated by sequencing (Eton Bioscience).

### AAV production and purification

HEK293T cells (ATCC CRL-11268) were obtained from the Duke Cell Culture Facility, which tests for mycoplasma contamination. Cells were maintained in culture medium containing DMEM medium (Gibco), 10% fetal bovine serum (Sigma F4135), and 1% penicillin-streptomycin in an incubator at 37°C and 5% CO_2_. Large-scale, high-titer viruses were produced in the Soderling lab using iodixanol (OptiPrep; Sigma) gradients as previously described (Uezu et al., 2016). Briefly, 1.5×10^7^ HEK293T cells were seeded onto each of six 15cm dishes per virus on the day before transfection. Cells were transfected using polyethylenimine (PEI MAX; Polysciences 24765-1) with 30μg helper plasmid pAdΔF6, 15μg serotype plasmid AAV2/9, and 15μg pAAV plasmid carrying the transgene. Cells were harvested 72 hours after transfection, resuspended in cell lysis buffer (15mM NaCl, 5mM T ris-HCl, pH 8.5), and subjected to 3 freeze-thaw cycles. The cell lysate was treated with 50U/ml benzonase, applied over an iodixanol density gradient (15%, 25%, 40%, and 60%), and ultracentrifuged for 2 hours at 60,000rpm in a Beckman Ti-70 rotor. The AAV-containing fraction was collected and concentrated by repeated washes with sterile PBS through a 100kDa filter (Amicon). The final volume of ∼200μ! was aliquoted and stored at −80°C until use. AAVs were titered using quantitative real-time PCR with primers against the ITR element (FWD: 5’GGAACCCCTAGTGATGGAGTT3’ and REV: 5’CGGCCTCAGTGAGCGA3’) (Aurnhammer et al., 2012).

Small-scale viruses were produced in the Soderling lab as previously described (Gao et al., 2019). Briefly, 2.5×10^5^ HEK293T cells were seeded onto one well in a 12-well plate per virus on the day before transfection. Cells in each well were transfected using polyethylenimine (PEI MAX; Polysciences 24765-1) with 0μg helper plasmid pAdΔF6, 0μg serotype plasmid AAV2/1, and 0μg pAAV plasmid carrying the transgene. Media was then changed to glutamine-free DMEM (ThermoFisher 11960044) supplemented with 1% GlutaMAX (Gibco) and 10% FBS (Sigma F4135). The AAV-containing supernatant medium was collected 72 hours after transfection and filtered through a 0.45μm Spin-X centrifuge tube filter (MilliporeSigma CLS8162). Small-scale viruses were stored at 4°C for up to one month until use.

### Immunocytochemistry and immunohistochemistry

For immunocytochemistry, cultured neurons were fixed at indicated timepoints with 4% PFA, 4% sucrose in PBS for 15min at 4°C. They were permeabilized with 0.25% Triton X-100 in PBS for 7min at room temperature and then blocked with blocking buffer containing 5% normal goat serum, 0.2% Triton X-100 in PBS for 1 hour at room temperature. Primary antibodies were diluted in blocking buffer and applied for 2 hours at room temperature. Coverslips were washed three times with 0.1% Triton X-100 in PBS for 5min at room temperature. Fluorescent secondary antibodies were diluted in blocking buffer and applied for 1 hour at room temperature, followed by counterstaining with DAPI. The coverslips were washed again and then mounted onto glass slides with FluorSave Reagent (Millipore 345789).

For immunohistochemistry, organotypic slices were cut at indicated timepoints from membranes with a scalpel and treated as free-floating sections. They were fixed with 4% PFA in PBS for 30min at 4°C and permeabilized with 1% Triton X-100 in PBS overnight at 4°C. They were then blocked in blocking buffer containing 5% normal goat serum, 0.1% Triton X-100, 0.03% NaNa_3_ in PBS for 6.5 hours at room temperature. Primary antibodies were diluted in blocking buffer and applied for 2 days at 4°C. Slices were washed three times with 0.2% Triton X-100 in PBS for 1 hour at room temperature. Fluorescent secondary antibodies were diluted in blocking buffer and applied overnight at 4°C, followed by counterstaining with DAPI. Slices were washed again and then mounted onto glass slides with FluorSave Reagent (Millipore 345789).

The following antibodies were used, with dilutions in blocking buffer indicated in parentheses. Primary antibodies: rat anti-HA (Roche 11867431001, 1:500), mouse anti-HA (Biolegend 901501, 1:500), mouse anti-V5 (ThermoFisher R960-25, 1:500), mouse anti-Myc (Santa Cruz sc-40, 1:250), mouse anti-bassoon (Abcam ab82958, 1:400), chicken anti-GFP (Abcam ab13970, 1:500), rabbit anti-RFP (Rockland 600-401-379, 1:500), rat anti-RFP (Chromotek 5F8, 1:500), rabbit anti-Homer1 (Synaptic Systems 160002, 1:500), guinea pig anti-Synapsin1 (Synaptic Systems 106104, 1:500), guinea pig anti-Vgat (Synaptic Systems 131004, 1:500), mouse anti-Gephyrin (Synaptic Systems 147011, 1:300), and mouse anti-Rac1 (BD Biosciences 610650, 1:250). Fluorophore-conjugated secondary antibodies: goat anti-chicken Alexa Fluor 488 (ThermoFisher A-11039, 1:500), goat anti-guinea pig Alexa Fluor 488 (ThermoFisher A-11073, 1:500), goat anti-guinea pig Alexa Fluor 647 (ThermoFisher A-21450, 1:500), goat anti-mouse Alexa Fluor 488 (ThermoFisher A-11029, 1:500), goat anti-mouse Alexa Fluor Plus 647 (ThermoFisher A-32728, 1:500), goat anti-rat Alexa Fluor 488 (ThermoFisher A-11006, 1:500), goat anti-rat Alexa Fluor 568 (ThermoFisher A-11077, 1:500), goat anti-rat Alexa Fluor 647 (ThermoFisher A-21247, 1:500), goat anti-rabbit Alexa Fluor 568 (ThermoFisher A-11036, 1:500), donkey anti-rabbit Alexa Fluor 647 (ThermoFisher A-31573, 1:500), and streptavidin Alexa Fluor 555 (ThermoFisher S-32355, 1:500).

### Validation of BioID probes

Hippocampal neuron cultures were prepared from *C57BL/6J* mice as described earlier. 1.5×10^6^ neurons were electroporated with μg pAAV-hSyn-BioID2-Linker-Synapsin1a-HA, pAAV-hSyn-BioID2-HA, or pCAG-GFP. 1.75×10^5^ WT and 1.65×10^4^ electroporated neurons were plated per well in a 24-well plate. Biotin (Sigma) was added to the media on DIV13 at a final concentration of 100μM, and neurons were fixed and stained on DIV14. Coverslips were imaged on a Zeiss LSM 710 inverted confocal microscope. All images were acquired by z-series (0.13μm intervals) using a 63x/1.4 numerical aperture (NA) oil-immersion objective. Maximum intensity projections from z-stacks along axons were analyzed in FIJI / ImageJ (Schindelin et al., 2012; Schneider et al., 2012). Intensity for HA (or GFP), streptavidin, and Bassoon was measured in both presynaptic terminals and the neighboring axonal shaft in small circular regions of interest (0.25μm diameter). Presynaptic terminals were identified as bouton-like swellings that colocalized with Bassoon.

Presynaptic enrichment was calculated by dividing the background-subtracted intensity in presynaptic terminals by the background-subtracted intensity in the corresponding axon. Localization values were then normalized to the average presynaptic enrichment of GFP, and biotinylation values were normalized to the average presynaptic enrichment of streptavidin in neurons expressing BioID2. Values for each axon were the average of at least four presynaptic terminals. All images were analyzed blinded to the condition. All probes were tested in at least three independent cultures.

### Synapsin *in vivo* BioID (iBioID)

P0-P1 *C57BL/6J* pups were anesthetized by hypothermia and intracranially injected with viruses as described previously (Uezu et al., 2016). AAV2/9-hSyn-BioID2-HA or AAV2/9-hSyn-BioID2-Linker-Synapsin1a-HA were bilaterally injected into the brain with a 10μ! Hamilton syringe (titer ∼3×10^13^ GC/ml; 0.8μ! per hemisphere), directed predominately into the hippocampus and cortex. Pups recovered on home cage bedding under a heat lamp and were returned to the dam together as a litter. From P21-P27, pups received daily subcutaneous injections of 24mg/kg biotin to increase biotinylation efficiency. At P28, brains were harvested from mice after deep isoflurane anesthesia. Cortices and hippocampi were quickly dissected, flash frozen in liquid nitrogen, and stored in a liquid nitrogen tank until ready for biotinylated protein purification.

A total of three independent purifications were performed. For each round of purification, the cortices and hippocampi of five mice were used per probe. First, synaptosomes were prepared from each mouse sample using a sucrose gradient. Frozen brain tissue was dounce homogenized in homogenization buffer (20mM sucrose, 5mM HEPES, 1mM EGTA, pH 7.4). Homogenate was centrifuged for 10min at 1,000 x *g* at 4°C. The supernatant (S1; crude cytosolic fraction) was transferred to a new tube and centrifuged for 20min at 12,000 x g at 4°C. The pellet (P2; crude synaptosomal fraction) was resuspended in resuspension buffer (320mM sucrose, 5mM Tris/Cl, pH 8.1), applied over a sucrose density gradient (1.2M, 1M, and 0.8M), and ultracentrifuged for 2 hours at 85,000 x *g* at 4°C in a Beckman SW 41 Ti rotor. All solutions contained a cocktail of protease and phosphatase inhibitors with final concentrations of 2μg/ml leupeptin, 2μg/ml pepstatin A, 1mM AEBSF, and 143μM sodium orthovanadate.

The purified synaptosomal fraction was carefully collected, and all synaptosomes expressing the same BioID probe were combined. Synaptosomes were lysed in RIPA buffer with sonication, followed by the addition of SDS to a final concentration of 1%. The lysate was then boiled for 5min by incubation in a 100°C water bath. After cooling on ice, samples were pre-cleared by the addition of Protein A agarose resin (Pierce) and rotation for 30min at 4°C. Beads were pelleted by centrifugation for 1min at 3,000 x *g* at 4°C, and the supernatant was collected with a 30g needle. To pulldown biotinylated proteins, high capacity NeutrAvidin agarose resin (Pierce) was added to the pre-cleared supernatant and rotated overnight for 14.5 hours at 4°C. Beads were pelleted by centrifugation for 1min at 3,000 x *g* at 4°C, and the supernatant was carefully removed using a 30g needle. Beads were extensively washed 2 times with 2% SDS, 2 times with 1% Triton X-100/1% deoxycholate/25mM LiCl, 2 times with 1M NaCl, and 5 times with 50mM ammonium bicarbonate in mass spectrometry-grade water (Honeywell). Biotinylated proteins were eluted into elution buffer (5mM biotin, 4% SDS, 20% glycerol, 10% beta-mercaptoethanol, 125mM Tris, pH 6.8 in mass spectrometry-grade water) by incubation for 5min in a 95°C heat block with periodic vortexing. Beads were pelleted by centrifugation for 1min @ 3,000 x g. The supernatant with eluted biotinylated proteins was carefully transferred to a low-protein-binding tube (Eppendorf) with a 30g needle and stored at −80°C.

### Quantitative mass spectrometry

The Duke Proteomics Core Facility received six eluents from streptavidin resins. Samples were supplemented with 10μ! 10% SDS, then reduced with 10mM dithiolthreitol for 30min at 80°C, alkylated with 20mM iodoacetamide for 45min at room temperature, and supplemented with a final concentration of 1.2% phosphoric acid and 384μ! of S-Trap (Protifi) binding buffer (90% MeOH/IOOmM TEAB). Proteins were trapped on the S-Trap, digested using 20ngμl sequencing grade trypsin (Promega) for 1 hour at 47°C, and eluted using 50mM TEAB, followed by 0.2% FA, and lastly using 50% ACN/0.2% FA. All samples were then lyophilized to dryness and resuspended in 12μ! 1%TFA/2% acetonitrile containing 25 fmol/μ! yeast alcohol dehydrogenase (ADH_YEAST). From each sample, 3μ! was removed to create a QC Pool sample which was run periodically throughout the acquisition period.

Quantitative ultraperformance liquid chromatography-tandem mass spectrometry (UPLC-MS/MS) was performed on 2.4μ! (∼20%) of each sample, using a nanoAcquity UPLC system (Waters Corp) coupled to a Thermo Orbitrap Fusion Lumos high resolution accurate mass tandem mass spectrometer (Thermo) via a nanoelectrospray ionization source. Briefly, the sample was first trapped on a Symmetry C18 20mm χ 180μm trapping column μl/min at 99.9/0.1 v/v water/acetonitrile), after which the analytical separation was performed using a 1μm Acquity HSS T3 C18 75μm χ 250mm column (Waters Corp) with a 90-min linear gradient of 5 to 40% acetonitrile with 0.1% formic acid at a flow rate of 400 nanoliters/minute (nl/min) with a column temperature of 55°C. Data collection on the Lumos mass spectrometer was performed in a data-dependent acquisition (DDA) mode of acquisition with a r=120,000 (@ m/z 200) full MS scan from m/z 375 - 1500 with a target AGC value of 2e5 ions followed by 30 MS/MS scans at r=15,000 (@ m/z 200) at a target AGC value of 5e4 ions and 45ms. A 20s dynamic exclusion was employed to increase depth of coverage. The total analysis cycle time for each sample injection was approximately 2 hours. The QC Pool was analyzed at the beginning, after every 3rd sample, and end of the sample set (3 times total). Individual samples were analyzed in a random order.

Following 9 total UPLC-MS/MS analyses (excluding conditioning runs, but including 3 replicate QC injections), data was imported into Proteome Discoverer 2.2 (Thermo Scientific Inc.), and analyses were aligned based on the accurate mass and retention time of detected ions (“features”) using Minora Feature Detector algorithm in Proteome Discoverer. Relative peptide abundance was calculated based on area-under-the-curve (AUC) of the selected ion chromatograms of the aligned features across all runs. The MS/MS data was searched against the SwissProt *M. musculus* database (downloaded in August 2017) with additional proteins, including yeast ADH1, bovine serum albumin, as well as an equal number of reversed-sequence “decoys” for false discovery rate determination. Mascot Distiller and Mascot Server (v2.5, Matrix Sciences) were utilized to produce fragment ion spectra and to perform the database searches. Precursor and product mass tolerances were set to 5ppm and 0.8Da, respectively, with full trypsin specificity and up to two missed cleavages. Database search parameters included fixed modification on Cys (carbamidomethyl) and variable modifications on Meth (oxidation) and Asn and Gln (deamidation). The overall dataset had 65,397 peptide spectral matches. Additionally, 286,563 MS/MS spectra were acquired for peptide sequencing by database searching. The data was annotated at a 1% peptide false discovery rate, resulting in identification of 5,406 peptides and 518 proteins.

### Differential protein expression and network analysis of proteomics data

Protein expression levels were intensity-scaled to the endogenously biotinylated proteins, pyruvate carboxylase (Q05920) and propionyl-CoA carboxylase (Q91ZA3). Imputation of missing values was performed after normalization (Karpievitch et al., 2012) using the MinDet method (Lazar et al., 2016). Missing values were replaced by the minimum value observed in each sample. For proteins found exclusively in BioID2-Synapsin but only in 2 out of the 3 replicates, the missing value was replaced by the average of the 2 replicates. To identify proteins specific to BioID2-Synapsin compared to BioID2, two-tailed t-tests were performed on log2-transformed protein intensities. P-values were corrected for multiple hypothesis testing using the FDR method. Fold changes were calculated by dividing the average protein intensity in BioID2-Synapsin by that in BioID2. To generate a high confidence list of hits, carboxylases, keratins, and other contaminants were removed as likely artifacts of overexpression or endogenously biotinylated proteins. These included proteins known to reside in other subcellular localizations such as the Golgi, endoplasmic reticulum, lysosome, nucleus, proteasome, and mitochondria, and those identified by PSD95-BirA (Uezu et al., 2016). To consider something specific for BioID2-Synapsin, we required at least 2 peptides to be identified in at least 2 replicates, with fold change greater than 32.5 over the negative control (BioID2) and adjusted p-value < 0.05.

Network figures were created using Cytoscape (v3.6) with node labels corresponding to the gene name for the identified protein. A non-redundant list of protein-protein interactions was assembled from the HitPredict database using the R package getPPIs (http://github.com/twesleyb/getPPIs), with additional hand annotation based on literature review. In all networks, node size is proportional to fold enrichment over BioID2 alone, and node shading corresponds to adjusted p-value. Clustergrams were based on gene set enrichment analysis using DAVID (https://david.ncifcrf.gov) (Dennis et al., 2003), as well as manual inspection based on UniProt database annotation and literature review. Neurological disease annotations were compiled based on UniProt, OMIM, and SFARI databases.

### HiUGE tagging of candidate genes

23 candidate genes were selected from the Synapsin iBioID proteome that had not previously been shown to localize to presynaptic terminals. These encoded for mostly actin regulators and 2 proteins of unknown function (see Table S1). Known protein isoforms, domains, binding regions, and localization signals were carefully assessed to minimize disruptions to protein function or localization by the insertion of a tag, and genes were generally tagged as close to the stop codon as reasonable. Mouse exon sequences were retrieved from the Ensembl genome browser (http://useast.ensembl.org) (Zerbino et al., 2018), and PAM sites (NGG) in these regions were identified using the CRISPOR guide RNA selection tool (http://crispor.tefor.net) (Haeussler et al., 2016). Target sequences were chosen for each gene based on specificity, predicted efficiency, and proximity to the stop codon.

Candidate guide RNAs were cloned as previously described (Gao et al., 2019). Briefly, oligos containing the 20bp target sequences with SapI overhangs were annealed. For some of the guides, an extra G was added at the start of the target sequence to enhance transcription under the U6 promotor. A combined restriction digestion and ligation reaction was performed to insert the annealed oligos behind the U6 promoter of the gene-specific GS-gRNA vector using SapI (NEB) and T4 DNA ligase (NEB). Correct integration of all oligos was confirmed by sequencing (Eton Bioscience). The 2xHA-V5-Myc HiUGE donor vector was created in all three open reading frames (ORFs) by inserting the payload sequence into the XbaI/PmlI sites of the HiUGE donor vector. The payload harbors a tandem array of six epitope tags (2x HA-, V5-, and Myc-tag), each separated by a spacing linker A(EAAAK)_2_A (Arai et al., 2001; Zhao et al., 2008). This design enables binding access of different epitope tag antibodies for flexible and synergistic labeling of modified endogenous proteins. Small-scale AAVs were prepared as described earlier for all candidate guides, 2xHA-V5-Myc HiUGE donors in the corresponding ORFs, and pAAV-Ef1a-GFP (as control).

To tag candidate genes, hippocampal neuron cultures were prepared from *H11Cas9* mice as described earlier. Neurons were plated densely, with dissociated cells from the hippocampi of 4 animals spread evenly across each 24-well plate. Small-scale AAVs (200ul each of guide and donor, or Ef1a-GFP alone) were added to each well on DIV0. As an additional negative control, neurons were also infected with only the donor AAV. Neurons were fixed and stained on DIV12-14 and then imaged on a Leica TCS SP8 inverted confocal microscope. Coverslips were first comprehensively viewed under the eyepieces to assess labeling efficiency, signal strength, and localization consistency. Candidate guides were not imaged further if there were no positive cells across 3 coverslips, or if the signal in positive cells was barely detectable above background fluorescence (see Table S1).

Images of whole neurons were acquired using a 20x/0.75 NA multi-immersion objective, and all images of axons were acquired by z-series (0.13μm intervals) using a 40x/1.3 NA oil-immersion objective. The sparse labeling of cells aided in the identification of axons, which were located as thin protrusions extending away from cell bodies for long distances with local swellings characteristic of presynaptic boutons. Maximum intensity projections from z-stacks along axons were analyzed in FIJI / ImageJ. Intensity for HA-V5-Myc (or GFP) and Synapsin1 was measured in both presynaptic terminals and the neighboring axonal shaft in small circular regions of interest (0.25μm diameter). Presynaptic terminals were identified as bouton-like swellings that colocalized with Synapsin1. Presynaptic enrichment was calculated by dividing the background-subtracted intensity in presynaptic terminals by the background-subtracted intensity in the corresponding axon. Enrichment values were then normalized to the average presynaptic enrichment of GFP. Values for each axon were the average of at least three presynaptic terminals, and axons from at least 5 neurons per guide were analyzed. All images were analyzed blinded to the condition. All guides were tested across four independent cultures. For display purposes, Synapsin1 puncta within axons were obtained by masking Synapsin1 fluorescence with a thresholded image of axonal HA-V5-Myc (or GFP). This was then merged with the original HA-V5-Myc (or GFP) image.

### Overrepresentation analysis for comparison of Arp2/3 and formin regulation

The enrichment of proteins involved in Arp2/3 or formin regulation in the Synapsin iBioID network was calculated using overrepresentation analysis (*Boyle et al., 2004*; *Rivals et al., 2007*) and hand-annotation based on literature review. The 31 genes involved in Arp2/3 regulation were: ActR2, ActR3, ArpC1, ArpC2, ArpC3, ArpC4, ArpC5, Cttn, Ctnnbp2, Cttnbp2nl, Cyfip1, Cyfip2, Abi1, Abi2, Brk1, Nckapl, Wasfl, Wasl, Wipf1, Wipf2, Wipf3, Wash1, WashC2, WashC3, WashC4, WashC5, Rac1, Cdc42, Abl1, Abl2, and Srgap3. The 36 genes involved in formin regulation were: Diaph1, Diaph2, Diaph3, Daam1, Daam2, Fmnll, Fmnl2, Fmnl3, Inf2, Fhdcl, Fhodl, Fhod3, Grid2ip, Fmn1, Fmn2, RhoA, RhoB, RhoC, RhoD, RhoF, Rac1, Cdc42, Fnbpl, Fnbpll, Rockl, Rock2, Dvl1, Dvl2, Dvl3, Baiap2, Pax6, Nckipsd, Src, Spire1, Srgap2, and Iqgap1. The enrichments of these genes in the Synapsin iBioID network was compared against a background of the entire mouse genome using a statistical test based on the hypergeometric distribution. A p-value, corresponding to the probability of obtaining by chance a number of annotated proteins equal or greater than the observed, was calculated using a custom script in MATLAB (MathWorks) implementing the equation:

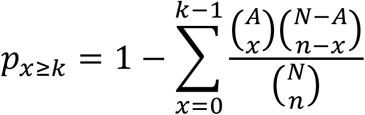

where N is the total number of genes in the background, A is the number of annotated genes in the background, n is the total number of genes in the sublist, and k is the number of annotated genes in the sublist. p-values for Arp2/3 and formin regulation were adjusted for multiple hypothesis testing using the FDR method.

### Pre-embedding immunogold electron microscopy

*C57BL/6J* mice were deeply anesthetized with pentobarbital (60mg/kg, i.p.) and then transcardially perfused with 0.9% NaCl followed by a mixture of 4% PFA and 0.1% glutaraldehyde (Electron Microscopy Sciences) in 0.1M phosphate buffer (PB), pH 7.4. Brains were removed and post-fixed overnight in 4% PFA without glutaraldehyde at 4°C. 60μm coronal sections from hippocampal CA1 were cut with a Leica VT1000 vibratome and processed for pre-embedding immunoelectron microscopy. Sections were incubated in primary antibodies diluted in 2% NDS. Primary antibodies used were as follows: mouse anti-Rac1 (BD Biosciences 610650, 1:100) and rabbit anti-ArpC2 (Millipore 07-227, 1:200).

Floating sections were treated for 30min in 1 % sodium borohydride in 0.1M PB to quench free aldehyde groups. The sections were incubated in 20% NDS for 30min to suppress nonspecific binding and then incubated for 12hr in the primary antibody, along with 2% NDS. Sections were incubated in biotinylated anti-rabbit IgG (Jackson) for 2hr. After rinses in PBS, sections were incubated in biotinylated donkey-anti rabbit or mouse IgG (Jackson) for 30 minutes, respective to the primary antibody. After washes in 0.1M PB, sections were incubated in 1.4nm Nanogold®-Streptavidin (Nanoprobes, 1:100) for 1hr at room temperature and rinsed in PB. Sections were washed in 0.1M Na acetate (to remove phosphate and chloride ions), followed by silver enhancement with IntensEM (GE Healthcare Life Sciences) or gold enhancement with GoldEnhance EM Plus (Nanoprobes) for approximately 8min.

Sections were processed as described above in control experiments, omitting primary antibody from the incubation solution. Sections were post-fixed in 0.5% osmium tetroxide in 0.1M PB for 30min. After dehydration in ascending ethanol series and contrasting with 1% uranyl acetate for 1hr in 70% EtOH, sections were incubated in propylene oxide and infiltrated with Durcupan resin (Sigma) and flat-mounted between sheets of Aclar (Electron Microscopy Sciences) within glass slides. Seventy-nanometer sections were cut, mounted on 300 mesh copper grids, contrasted with lead citrate (Ultrostain II; Leica), and examined in a JEOL TEM-1011 electron microscope at 80 kV; images were collected with a Megaview 12-bit 1024 χ 1024 CCD camera. Electron micrographs were taken from randomly selected fields, focusing on the middle one third of hippocampal CA1 *stratum radiatum.*

### Quantitative analysis of immunogold labeling and synaptic vesicle position

Synaptic vesicle distances, immunogold particle distances, and profile areas were measured from electron micrographs using ImageJ 1.52a. The “axo-dendritic” positions of immunogold particles were calculated as previously described (Racz and Weinberg, 2004). Briefly, we defined the lateral edges of the PSD for a random sample of clearly-defined synapses, and measured the shortest distance from the center of each gold particle to the outer layer of the presynaptic membrane. Normalized axodendritic position (dN, the fraction of the distance from the presynaptic plasma membrane) was computed according to the equation:

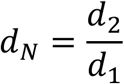

where d_2_ is the axodendritic distance from the presynaptic membrane to the particle or synaptic vesicle, and d_1_ is the distance of the axodendritic diameter of the given terminal profile. Thus, 0 corresponds to a particle at the presynaptic membrane of the terminal, while 1 corresponds to a particle at the opposite plasma membrane, at the furthest possible position on the terminal profile. For this measurement, only particles lying within the axon terminal were considered.

### Mixed hippocampal cultures for presynaptic isolation

Hippocampal neuron cultures were prepared from *C57BL/6J* (WT), *Rac1^fl/fl^,* or *ArpC3^fl/fl^;Ai14* mice as described earlier. For presynaptic Rac1 experiments, 1.75×10^6^ *Rac1^fl/fl^* neurons were electroporated with 1μg pAAV-hSyn-ChR2-EYFP, and 5×10^6^ WT neurons were electroporated with 3μg pBA-tdTomato. 5×10^5^ WT electroporated and 1.75×10^5^ *Rac1^fl/fl^* electroporated neurons were plated per well in a 24-well plate. This resulted in the sparse seeding of *Rac1^f/fl^;ChR2-EYFP* neurons, as ∼80% of electroporated neurons do not survive. On DIV10, AAV2/9-hSyn-Cre was added to half of wells (0.5μ! of 3.21×10^13^ GC/ml per well), with sterile PBS as loading control. For presynaptic Arp2/3 experiments, 1.5×10^6^ *ArpC3^f/fl^;Ai14* neurons were electroporated with μg pAAV-hSyn-Ch R2-EYFP or pBA-tdTomato. 1.75×10^5^ WT and 0.6×10^5^ *ArpC3^fl/fl^;Ai14* electroporated neurons were plated per well in a 24-well plate. On DIV10, AAV2/9-hSyn-Cre was added to half of wells (0.5μ! of 3.21×10^13^ GC/ml per well), with sterile PBS as loading control. For PA-Rac1 experiments, 1.5×10^6^ WT neurons were electroporated with 0μg pCAG-ChrimsonR-tdT, pCAG-ChrimsonR-tdT-P2A-PA Rac1 DN, or pCAG-ChrimsonR-tdT-P2A-PA Rac1 CA, and 1μg pCDNA3. 1.75×10^5^ WT and 1.5×10^5^ electroporated neurons were plated per well in a 24-well plate. For immunostaining of mixed cultures, neurons were fixed and stained on DIV16. Coverslips were imaged on a Zeiss LSM 710 inverted confocal microscope. All images were acquired by z-series (0.13μm intervals) using a 63x/1.4 numerical aperture (NA) oil-immersion objective.

### Electrophysiology

Somatic whole-cell currents were recorded from cultured hippocampal neurons on DIV16-18 under a Zeiss Axio Examiner.D1 upright microscope equipped with IR-DIC optics. Patch pipettes (4-7 MΩ) were created from borosilicate glass capillaries (Sutter Instrument) using a P-97 puller (Sutter Instrument). Coverslips were superfused with artificial CSF (aCSF) containing 124mM NaCl, 26mM NaHCO_3_, 10mM dextrose, 2mM CaCh, 3mM KCl, 1.3mM MgSO_4_, and 1.25mM NaH2PO4 (310 mOsm/L), continuously bubbled at room temperature with 95% O_2_ and 5% CO_2_. For voltage-clamp experiments, pipette intracellular solution contained 135mM Cs-methanesulfonate, 8mM NaCl, 10mM HEPES, 0.3mM EGTA, 10mM Na_2_phosphocreatine, 4mM MgATP, 0.3mM Na_2_GTP, 5mM TEA-Cl, and 5mM QX-314 (pH 7.3 with CsOH, 295 mOsm/L). Light-evoked EPSCs were recorded at −70mV holding potentials in aCSF with the following modifications: 100μM picrotoxin, 10μM bicuculline methiodide, and 50μM D-AP5. Light-evoked IPSCs were recorded at 0mV holding potentials in aCSF with the following modifications: 50μM D-AP5 and 20μM CNQX. For strontium substitution experiments, 4mM SrCh replaced 2mM CaCl_2_ in aCSF. For current-clamp recordings, pipette intracellular solution contained 135mM K-methanesulfonate, 8mM NaCl, 10mM HEPES, 0.3mM EGTA, 4mM MgATP, and 0.3mM Na_2_GTP (pH 7.3 with KOH, 295 mOsm/L). Light-evoked action potentials were recorded at 0pA holding currents in aCSF with the following modifications: 20μM CNQX, 50μM D-AP5, 100μM picrotoxin, and 10μM bicuculline methiodide. No corrections were made for the 8.5-9.0mV estimated liquid junction potentials of these solutions. All drugs were purchased from MilliporeSigma or Tocris.

Light was delivered through the objective using an LED light source (CoolLED pE-300ultra) with 460nm and 525-660m excitation peaks and corresponding filter sets, with the shutter controlled by TTL inputs. 1ms pulses of 460nm light were used to activate ChR2, while 3ms pulses of 525-660nm light were used to activate ChrimsonR. Light intensities were kept constant across all recordings (20% for EPSCs and 10% for IPSCs). For paired pulse and strontium substitution experiments, neurons were stimulated at no more than 0.1Hz between sweeps. Recordings were not continued if the light-evoked current was not monosynaptic.

Series resistance was monitored throughout all voltage-clamp recordings with brief 5mV hyperpolarizing pulses, and only recordings which remained stable over the period of data collection were analyzed. Data were recorded with a Multiclamp 700B amplifier (Molecular Devices), digitized at 50kHz with a Digidata 1550 (Molecular Devices), and low-pass filtered at 1 kHz. For PA-Rac1 experiments, recordings were conducted in the dark with monitors and other light sources covered by blue light filters (135 Deep Golden Amber; Lee). These coverslips were allowed to recover for 15min in the dark after transferring them to the recording chamber, and between each recording. For all recordings, the experimenter was not blinded to the condition. All experiments were repeated on at least three independent cultures.

For voltage-clamp experiments, EPSC and IPSC amplitudes were manually detected and calculated offline using MiniAnalysis (Synaptosoft) with suggested detection parameters. Paired-pulse ratio (PPR) was calculated as the average of 6-10 trials conducted every 10s. Quantal events from strontium substitution experiments were also manually detected in MiniAnalysis with a threshold of 5pA. All events were counted in 500ms (for qEPSCs) or 1s (for qIPSCs) time windows after stimulation, with stimulation every 10s for 5min. For 20Hz stimulation trains, a linear regression was performed on the final 10 data points on cumulative current curves, as specified by the “train method” (Stevens and Williams, 2007; Thanawala and Regehr, 2016). The size of the readily releasable pool (RRP) was quantified as the y-intercept of the line, the synaptic vesicle replenishment rate as the slope of the line, and the initial release probability (p) as the amplitude of the first current divided by the RRP size. Analysis of charge transfer, kinetics, and basal current was done in Clampfit 10 (Molecular Devices). For current-clamp experiments, action potentials were counted if the peak was greater than 0mV. Action potential waveforms were also analyzed using Clampfit 10. All experiments were analyzed blinded to the condition.

### Quantification of axonal synapse density

The sparse seeding of *ArpC3^fl/fl^;Ai14;tdTomato* neurons allowed for the identification of long axonal processes away from cell bodies. Maximum intensity projections from z-stacks along these axons were analyzed in FIJI / ImageJ. Presynaptic (Synapsin1 or Vgat) puncta within axons were obtained by masking their fluorescence with a thresholded image of the tdTomato axonal fill. A custom Puncta Analyzer plugin for ImageJ 1.29 written by Barry Wark (Ippolito and Eroglu, 2010) was then used to calculate the number of presynaptic puncta within axons that was colocalized with postsynaptic puncta (Homer1 or Gephyrin) in the field. The length of each axon was determined in FIJI/ ImageJ using the Simple Neurite Tracer plugin (Longair et al., 2011). Axonal synapse density was calculated as the number of colocalized puncta divided by the length of the axon. All experiments were repeated on at least three independent cultures, and all images were analyzed blinded to the condition.

### Microinjection of organotypic hippocampal slices

Organotypic hippocampal slices were prepared in the Yasuda lab. Slices were microinjected in CA3 on DIV10-13 to induce expression of the Rac1 FLIM donor (AAV2/9-hSyn-mEGFP-Rac1) and acceptor (AAV2/9-hSyn-mCherry-PBD2-mCherry). Briefly, AAVs were mixed together in a 1:2 donor: acceptor ratio (final titer of each ∼1-2×10^12^ GC/ml) with 10% Fast Green FCF dye. Pipettes were created from glass capillaries (VWR) using a P-1000 puller (Sutter Instrument) and back-filled with AAV mixture. The mixture was microinjected into the pyramidal cell layer of CA3 using a Picospritzer III (Parker) set to 18psi with a pulse duration of 50ms, and then slices on culture inserts were returned to the incubator.

### Two-photon fluorescence lifetime imaging (2pFLIM)

On DIV17-24, at least 7 days after microinjection, 2pFLIM was conducted on synaptic boutons in CA1. Organotypic slices were cut from inserts using a scalpel and transferred to an imaging chamber. Slices were superfused with artificial CSF (aCSF) containing 124mM NaCl, 3mM KCl, 1.25mM NaH2PO4, 26mM NaHCO_3_, 10mM dextrose, 4mM CaCh, and 1.3mM MgSO_4_ (310 mOsm/L), continuously bubbled at room temperature with 95% O_2_ and 5% CO_2_. A concentric bipolar electrode (CBAPC75; FHC) was placed in the Schaffer collaterals and attached to an ISO-Flex stimulus isolator (AMPI). A recording electrode filled with aCSF was placed in CA1 *stratum radiatum,* and stimulation intensity was adjusted to evoke field potentials at half-maximum amplitude. Data were recorded with a Multiclamp 700B amplifier (Molecular Devices) interfacing with custom software. Slices with mistargeting of viral microinjections or evidence of epileptiform activity were discarded. For pharmacological experiments, 0μM TTX or 300μM CdCh was washed onto slices after evoking field potentials, and slices were then incubated with the compound for at least 30min before imaging.

2pFLIM using a custom-built microscope was performed as previously described (Murakoshi et al., 2011). GFP and mCherry were excited with a Ti-sapphire laser (Chameleon; Coherent) tuned to a wavelength of 920nm. All samples were imaged using <2mW laser power measured below the objective. Fluorescence was collected using a 60x/1.0 NA water-immersion objective (Olympus), divided with a dichroic mirror (565nm) and detected with two separate photoelectron multiplier tubes (PMTs) placed downstream of two wavelength filters (510/70-2p for green and 620/90-2p for red; Chroma). PMTs with low transfer time spread (H7422-40p; Hamamatsu) were used for both red and green channels. Photon counting for fluorescence lifetime imaging was performed using a time-correlated single photon counting board (SPC-150; Becker and Hickl) controlled with custom software, while fluorescence images were acquired using a separate data acquisition board (PCI-6110; National Instrument). 2pFLIM images were collected with 64 χ 64 pixels at 128 ms/frame, with 80 frames per image. A new image was taken every 10s over a period of 5min. 2s stimulation at 50Hz was initiated by hand after a 2min baseline period. All conditions were imaged over at least four independent slices.

To measure the fraction of donor bound to acceptor, we fit a fluorescence lifetime curve summing all pixels over a whole image with a double exponential function convolved with the Gaussian pulse response function:

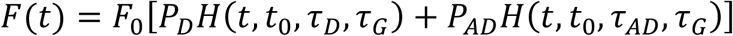

where *τ_*AD*_* is the fluorescence lifetime of donor bound with acceptor, *P_D_* and *P_AD_* are the fraction of free donor and donor bound with acceptor, respectively, and *H(t)* is a fluorescence lifetime curve with a single exponential function convolved with the Gaussian pulse response function:

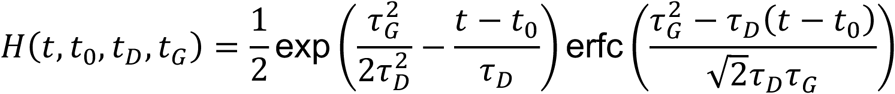

in which *τ_D_* is the fluorescence lifetime of the free donor, *τ_G_* is the width of the Gaussian pulse response function, *F_0_* is the peak fluorescence before convolution, *T_0_* is the time offset, and erfc is the error function.

We fixed *τ_D_* to the fluorescence lifetime obtained from free EGFP (2.6ns) and *τ_AD_* to 1.1ns based on previous experiments (Hedrick et al., 2016). To generate the fluorescence lifetime image, we calculated the mean photon arrival time, (t), in each pixel as:

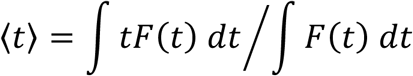

Then, the mean photon arrival time is related to the mean fluorescence lifetime, (*T*), by an offset arrival time, *T*_0_, which is obtained by fitting the whole image with the following equation:

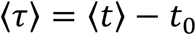

Finally, the binding fraction *(P_AD_)* was calculated for small regions of interest in presynaptic boutons as:

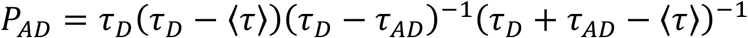

Change in binding fraction was calculated by subtracting the average value before stimulation. Data with lifetime fluctuations in the baseline that were greater than 0.1ns were excluded before further analysis. Lifetime drift was not corrected in the analysis.

### Quantification of presynaptic boutons containing synapsin

Maximum intensity projections from z-stacks along axons in CA1 of organotypic slices were analyzed in FIJI / ImageJ. Synapsin1 puncta within axons were obtained by masking their fluorescence with a thresholded image of axons expressing both mCherry-PBD2-mCherry and mEGFP-Rac1. Presynaptic boutons were manually marked as swellings along axons, and then Synapsin1 puncta were independently marked using the custom Puncta Analyzer plugin for ImageJ 1.29 (Ippolito and Eroglu, 2010). Swellings containing at least one Synapsin1 puncta were counted as Synapsin+ boutons. Images were analyzed from axons in three different slices.

### Statistics

For all graphs, center values represent mean, and error bars represent standard error of the mean (SEM). Details of exact sample sizes and statistical tests used can be found in figure legends. No statistical methods were used to predetermine sample sizes, but our sample sizes are similar to those reported in previous publications (Hedrick et al., 2016; Spence et al., 2019). Statistical analysis was performed in Prism 8 (GraphPad) and MATLAB R2017a (MathWorks). We compared independent sample means using two-tailed t-tests, one-way ANOVAs, two-way ANOVAs, and repeated measures ANOVAs as appropriate. ANOVAs were followed by Tukey’s, Dunnett’s, or Sidak’s multiple comparisons tests. When required, hypergeometric tests and t-tests were adjusted for multiple comparisons using the FDR method. We confirmed necessary parametric test assumptions using the Shapiro-Wilk test (normality). Violations in test assumption were corrected by transformations when possible; otherwise, the equivalent non-parametric tests were applied instead. Type-1 error rates for all tests were set at 0.05.

## DATA AVAILABILITY

All data generated in this study are included in the manuscript and supporting files. Raw proteomics data have been deposited to the ProteomeXchange Consortium via the PRIDE partner repository with the dataset identifier PXD019342 (Note to reviewers – please login with the reviewer account, username, password: XRNgjLCm).

---

**Figure 1—figure supplement 1.**
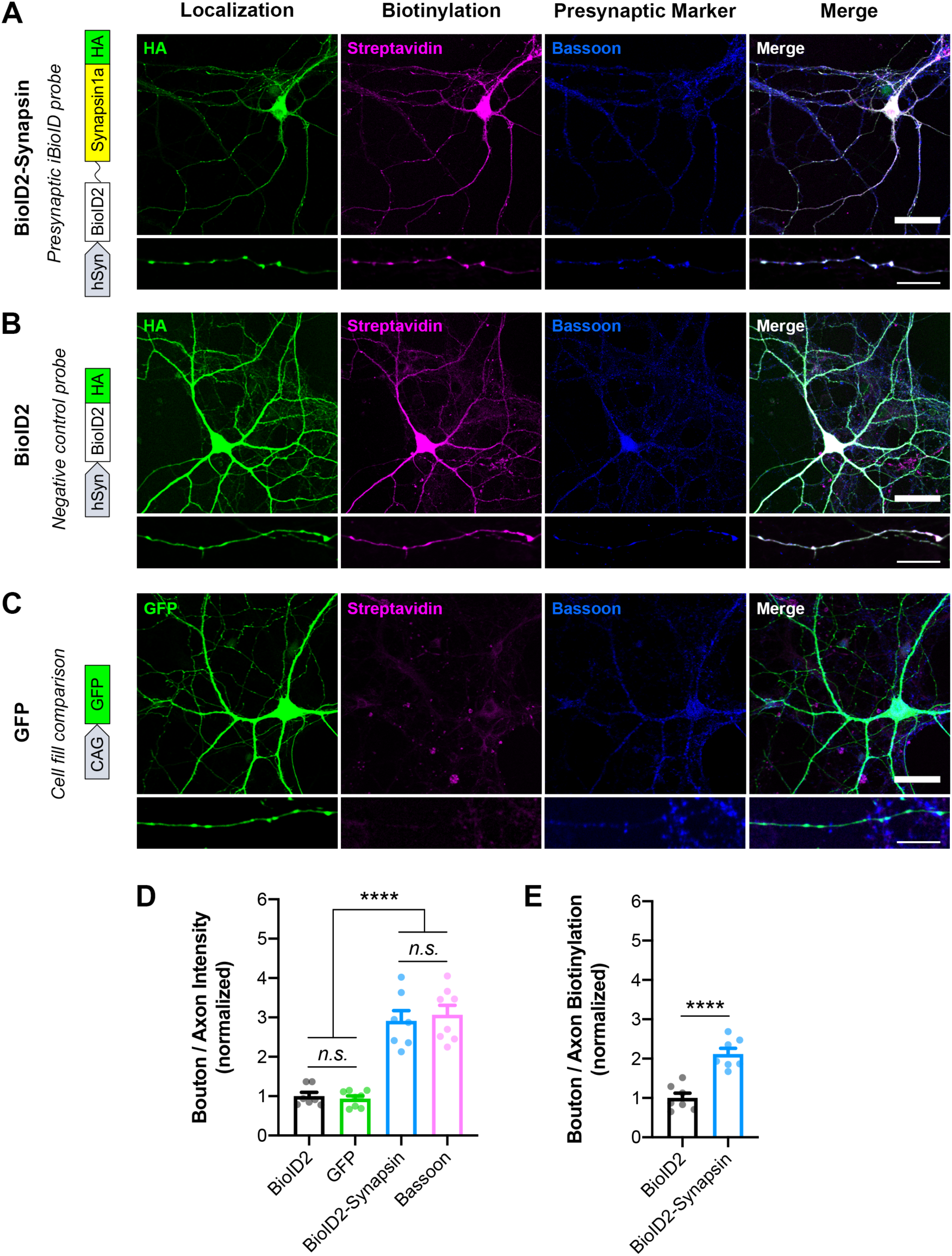
Validation of Synapsin iBioID probes in cultured hippocampal neurons. **(A-C)** Representative images of localization (HA or GFP; green), biotinylation (Streptavidin; magenta), and a presynaptic marker (Bassoon; blue) in neurons expressing **(A)** BioID2-Linker-Synapsin1a-HA, the iBioID bait, **(B)** BioID2-HA, the negative control, and **(C)** GFP, a soluble fill. Scale bars, 40μm. Insets show staining along axons. Scale bars, 10μm. **(D)** Presynaptic enrichment of localization for all probes (BioID2 n=7 neurons/3 cultures, GFP n=8/3, BioID2-Synapsin n=7/3, Bassoon n=8/3); one-way ANOVA (F3,26=60.18, p<0.0001) with Tukey’s multiple comparisons test: BioID2 vs GFP (p=0.9526), BioID2 vs BioID2-Synapsin (p<0.0001), BioID2 vs Bassoon (p<0.0001), GFP vs BioID2-Synapsin (p<0.0001), GFP vs Bassoon (p<0.0001), BioID2-Synapsin vs Bassoon (p=0.9655). **(E)** Presynaptic enrichment of biotinylation for iBioID probes (BioID2 n=7/3, BioID2-Synapsin n=7/3); t-test (t12=5.943, p<0.0001). All data are mean ± SEM. ****p<0.0001, *n.s*. not significant.

**Figure 3—figure supplement 1.**
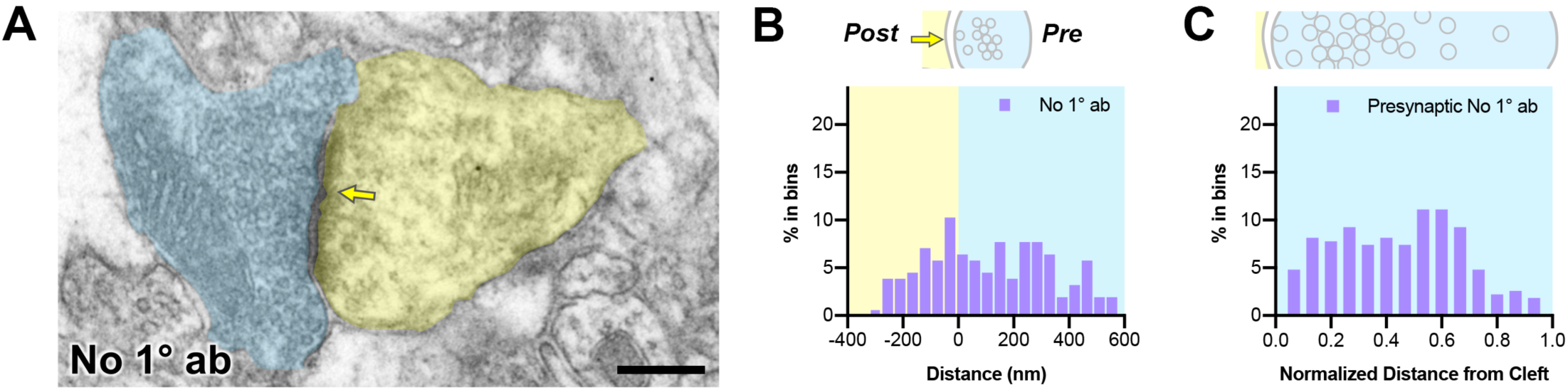
Negative control for immunogold electron microscopy. **(A)** Representative pre-embedding immunogold-labeled electron micrograph in mouse hippocampal CA1 with primary antibody omitted. Dendritic spines are pseudocolored yellow, presynaptic terminals are pseudocolored blue, and a yellow arrow points to the synaptic cleft. Scale bar, 200nm. **(B)** Axodendritic distribution of gold particles at the synapse in negative control samples (n=102 synapses). **(C)** Presynaptic distribution of gold particles in negative control samples. Distances were normalized from the synaptic cleft based on the axodendritic length of the presynaptic terminal.

**Figure 4—figure supplement 1.**
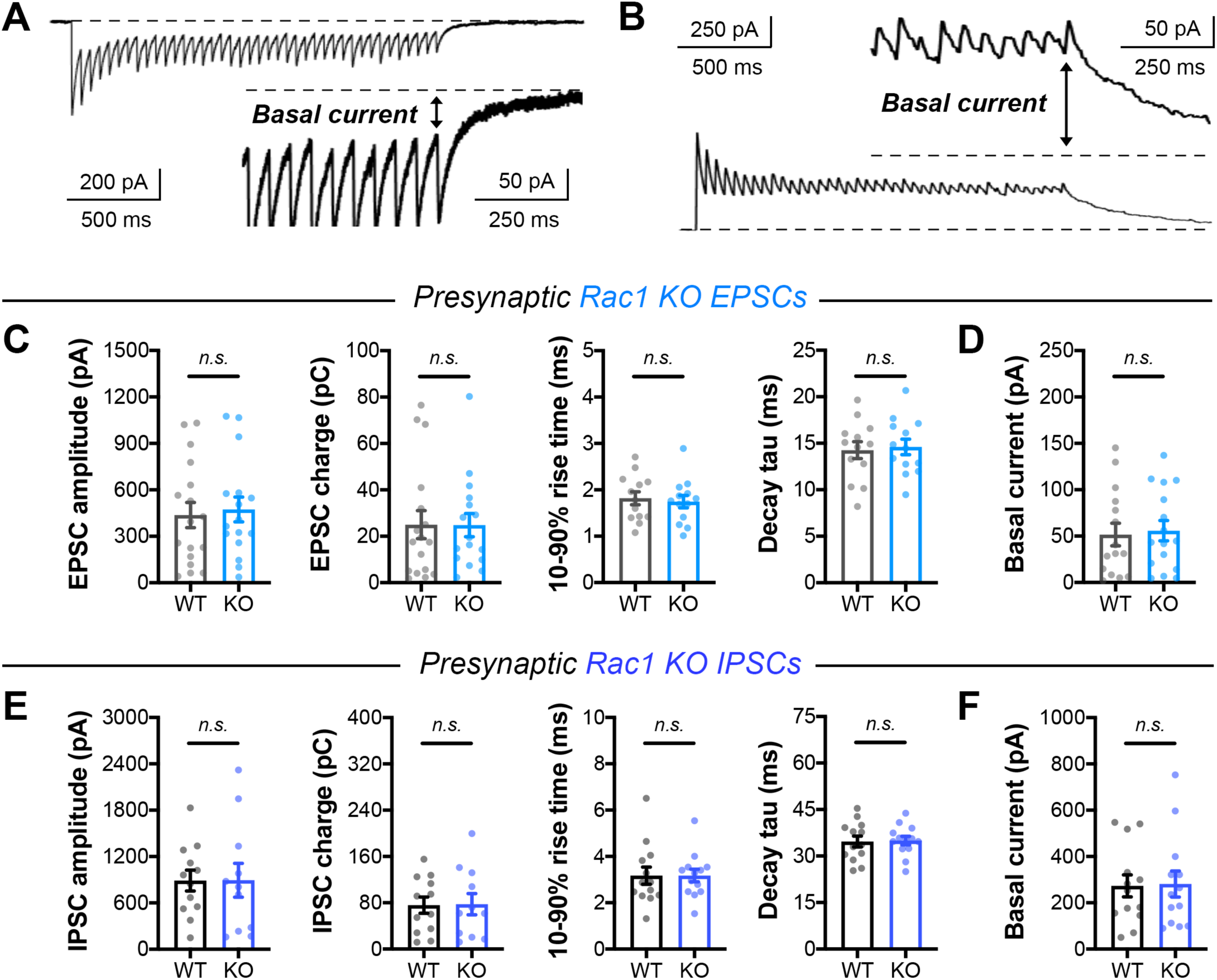
Single evoked currents and asynchronous release in *Rac1* neurons. **(A-B)** Visual representation of steady-state basal current in **(A)** EPSC trains and **(B)** IPSC trains. **(C)** Quantification of single evoked EPSCs in *Rac1* cultures (WT n=17 neurons/3 cultures, KO n=16/3) for amplitude (t-test, t31=0.3127, p=0.7566), charge (Mann-Whitney U-test, U=123, p=0.6567), rise time (t-test, t31=0.5223, p=0.6051), and decay time constant (t-test, t31=0.08846, p=0.9301). **(D)** Basal current in *Rac1* EPSC trains (WT n=15/3, KO n=16/3); t-test (t29=0.2560, p=0.7998). **(E)** Quantification of IPSCs in *Rac1* cultures (WT n=12/3, KO n=11/3) for amplitude (t-test, t21=0.01487, p=0.9883), charge (t-test, t21=0.07053, p=0.9444), rise time (t-test, t21=0.5311, p=0.6009), and decay time constant (t-test, t21=0.3887, p=0.7014). **(F)** Basal current in *Rac1* IPSC trains (WT n=13/3, KO n=13/3); Mann-Whitney U-test (U=83, p=0.9598). All data are mean ± SEM. *n.s*. not significant.

**Figure 5—figure supplement 1.**
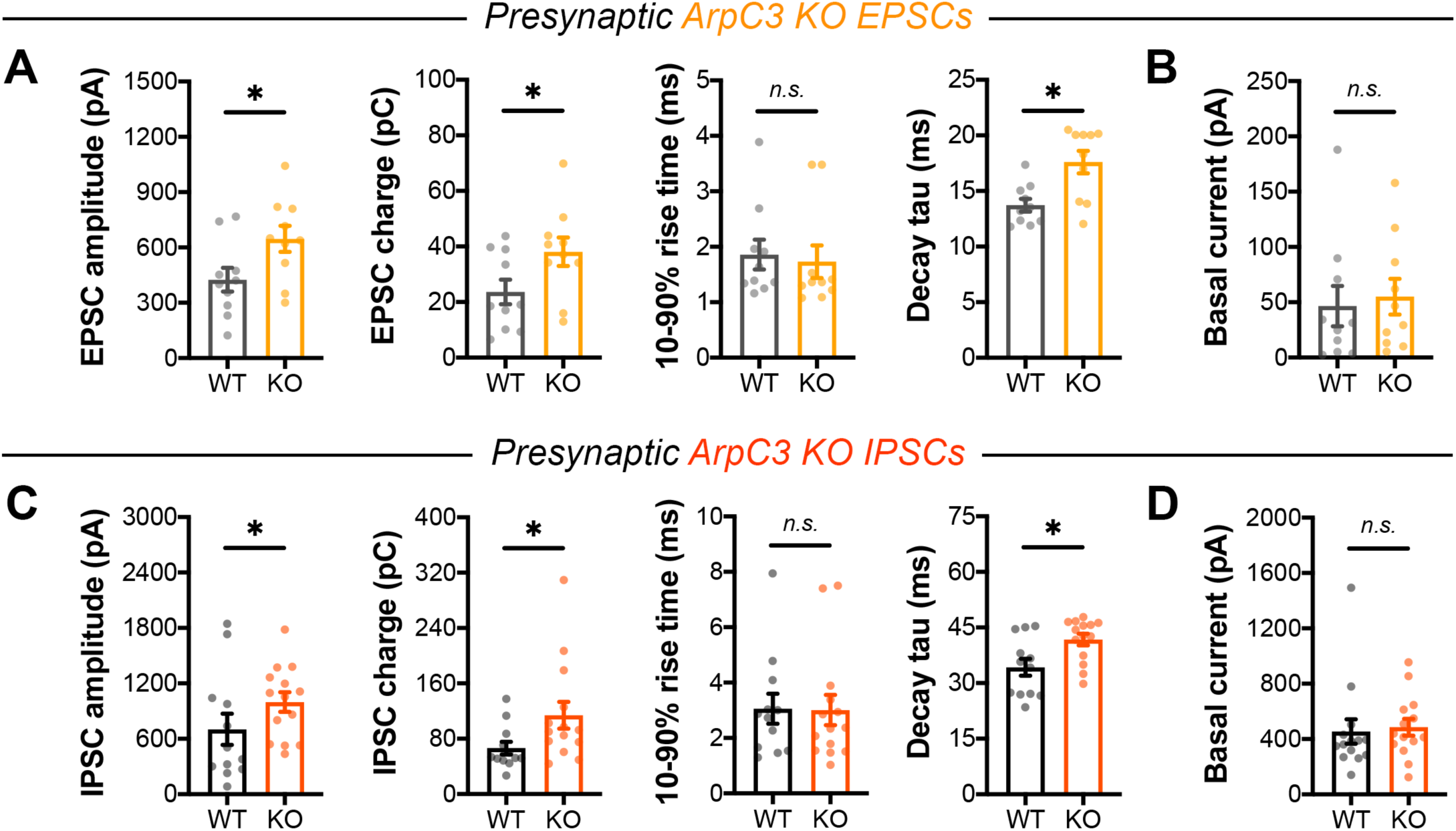
Single evoked currents and asynchronous release in *ArpC3* neurons. **(A)** Quantification of EPSCs in *ArpC3* cultures (WT n=10/3, KO n=10/3) for amplitude (t-test, t18=2.323, p=0.0321), charge (t-test, t18=2.127, p=0.0475), rise time (Mann-Whitney U-test, U=38, p=0.3811), and decay time constant (Mann-Whitney U-test, U=17, p=0.0111). **(B)** Basal current in *ArpC3* EPSC trains (WT n=10/3, KO n=10/3); Mann-Whitney U-test (U=42, p=0.5787). **(C)** Quantification of IPSCs in *ArpC3* cultures (WT n=12/3, KO n=14/3) for amplitude (Mann-Whitney U-test, U=44, p=0.0407), charge (Mann-Whitney U-test, U=42, p=0.0310), rise time (Mann-Whitney U-test, U=78, p=0.7810), and decay time constant (Mann-Whitney U-test, U=36, p=0.0127). **(D)** Basal current in *ArpC3* IPSC trains (WT n=14/3, KO n=14/3); Mann-Whitney U-test (U=72, p=0.2456). All data are mean ± SEM. *p<0.05, *n.s*. not significant.

**Figure 5—figure supplement 2.**
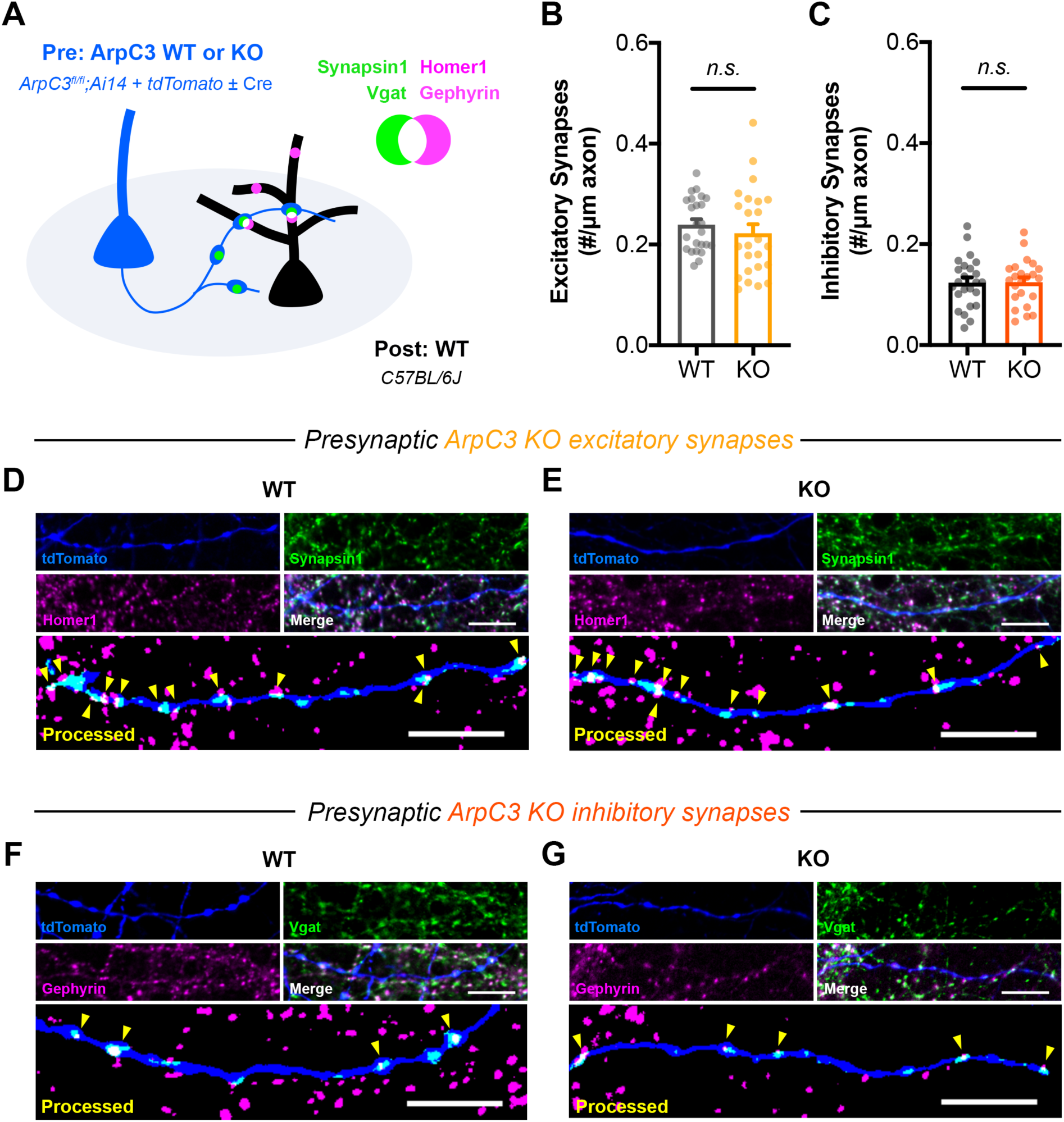
ArpC3 loss does not affect the density of synapses formed along axons. **(A)** Schematic of mixed hippocampal neuron cultures to isolate effects of *ArpC3* knockout on axonal synapse density. *ArpC3fl/fl;Ai14* neurons were electroporated with tdTomato and sparsely seeded amongst WT neurons on DIV0. To limit developmental effects, AAV-hSyn-Cre was added on DIV10 to half the coverslips. Neurons were fixed on DIV16 and stained for excitatory (Synapsin1, Homer1) or inhibitory (Vgat, Gephyrin) synapse markers. **(B)** Excitatory synapse density along axons (WT n=24 neurons/3 cultures, KO n=24/3); t-test (t46=0.8180, p=0.4176). **(C)** Inhibitory synapse density along axons (WT n=24/3, KO n=23/3); t-test (t45=0.9572, p=0.9572). **(D-E)** Representative images of **(D)** WT and **(E)** KO axons stained for tdTomato (blue), Synapsin1 (green), and Homer1 (magenta). Synapsin1 puncta were masked inside tdTomato+ axons and counted as synapses (yellow arrows) if they colocalized with Homer1 puncta. Scale bars: 10μm. **(F-G)** Representative images of **(F)** WT and **(G)** KO axons stained for tdTomato (blue), Vgat (green), and Gephyrin (magenta). Vgat puncta were masked inside tdTomato+ axons and counted as synapses (yellow arrows) if they colocalized with Gephyrin puncta. Scale bars, 10μm. All data are mean ± SEM. *n.s*. not significant.

**Figure 5—figure supplement 3.**
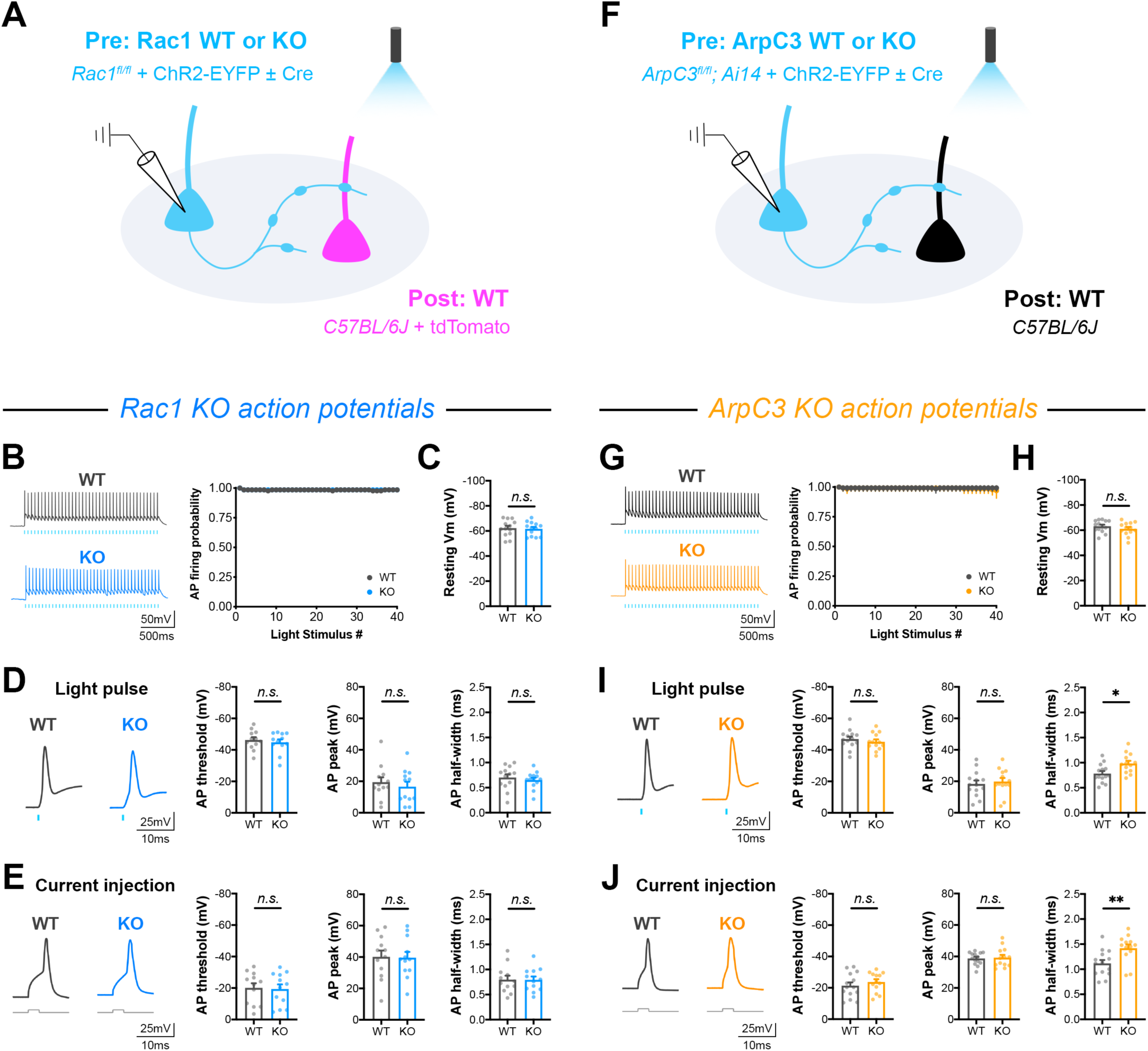
Action potential firing and intrinsic membrane properties in *Rac1* and *ArpC3* neurons. **(A)** Schematic of *Rac1* mixed hippocampal neuron cultures. Current clamp recordings were conducted from ChR2+ neurons with light delivered through the objective by a 460nm LED (WT n=12 neurons/3 cultures, KO n=12/3). **(B)** Probability of firing action potentials during a 20Hz light stimulation train (two-way repeated measures ANOVA, F1,22=0.004558, p=0.9468). **(C)** Resting membrane potential (t-test, t22=0.2579, p=0.7989). **(D)** Waveforms of light-evoked action potentials and quantification of threshold (Mann-Whitney U-test, U=65, p=0.7125), height (t-test, t22=0.6447, p=0.5258), and half-width (t-test, t22=0.8472, p=0.4060). **(E)** Waveforms of action potentials elicited by current injection and quantification of threshold (t-test, t22=0.1823, p=0.8571), height (t-test, t22=0.1056, p=0.9168), and half-width (t-test, t22=0.0502, p=0.9604). **(F)** Schematic of *ArpC3* mixed hippocampal neuron cultures and current clamp recordings (WT n=14/3, KO n=13/3). **(G)** Probability of firing action potentials during a 20Hz light stimulation train (two-way repeated measures ANOVA, F1,25=0.07845, p=0.7817). **(H)** Resting membrane potential (t-test, t25=1.068, p=0.2959). **(I)** Waveforms of light-evoked action potentials and quantification of threshold (t-test, t25=0.7780, p=0.4438), height (t-test, t25=0.4700, p=0.6424), and half-width (t-test, t25=2.745, p=0.0111). **(J)** Waveforms of action potentials elicited by current injection and quantification of threshold (t-test, t25=0.8628, p=0.3964), height (t-test, t25=0.2601, p=0.7969), and half-width (t-test, t25=2.991, p=0.0062). All data are mean ± SEM. *p<0.05, **p<0.01, *n.s*. not significant.

**Figure 7—figure supplement 1.**
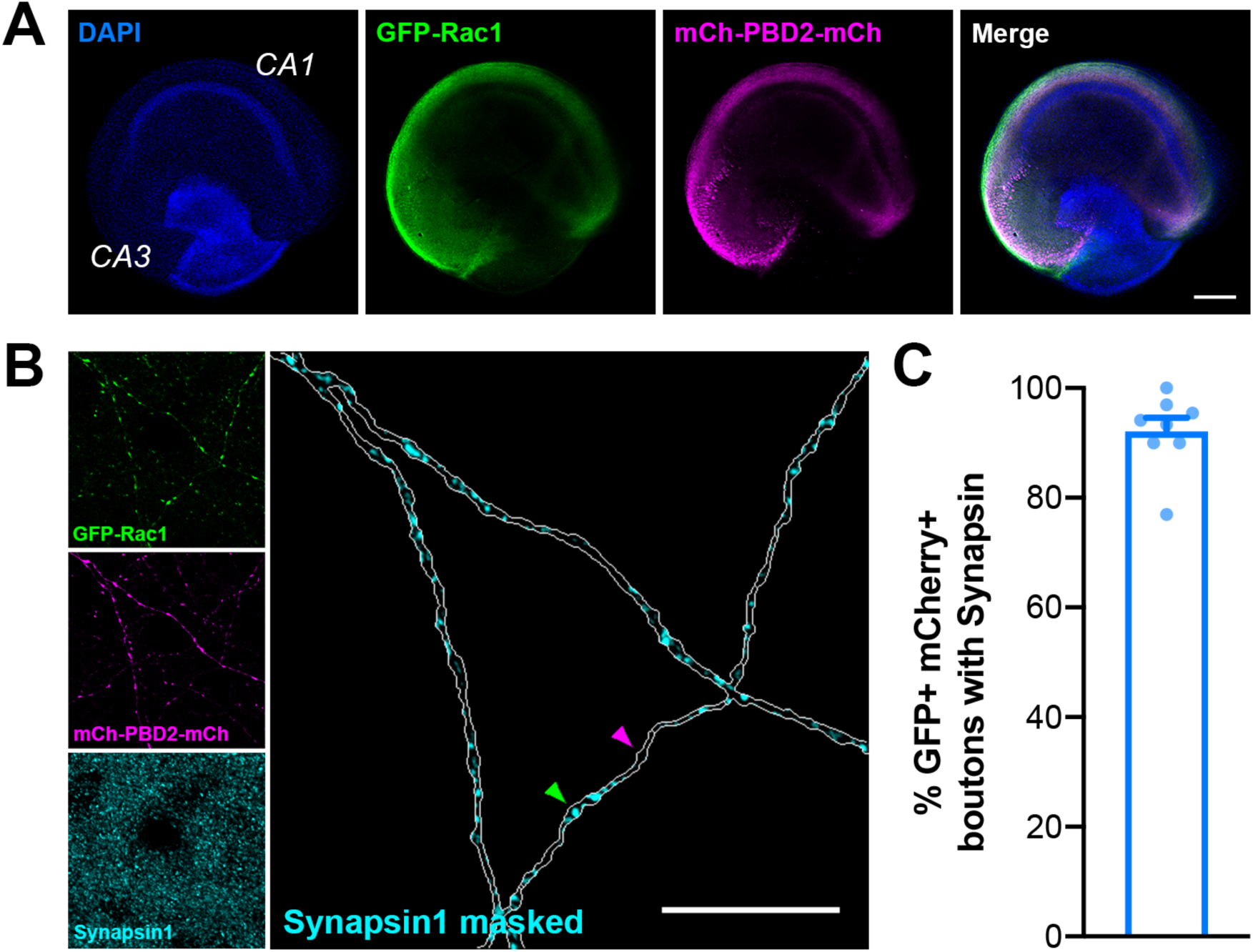
Presynaptic boutons in organotypic slices used for 2pFLIM contain synapsin. **(A)** Representative image of an organotypic hippocampal slice microinjected in CA3 with AAV-mEGFP-Rac1 and AAV-mCherry-PBD2-mCherry on DIV10 and fixed and stained on DIV20 for DAPI (blue), GFP (green), mCherry (magenta), and Synapsin1 (cyan). Scale bar, 300μm. **(B)** Representative image of axons in CA1 expressing the Rac1 activity sensor. Green arrow points to a Synapsin1+ bouton, and magenta arrow points to a bouton without Synapsin1 staining. Scale bar, 15μm. **(C)** Quantification of percent of GFP+ mCherry+ boutons with Synapsin1 staining (n=142 boutons/8 slices). All data are mean ± SEM.

**Table S1.**
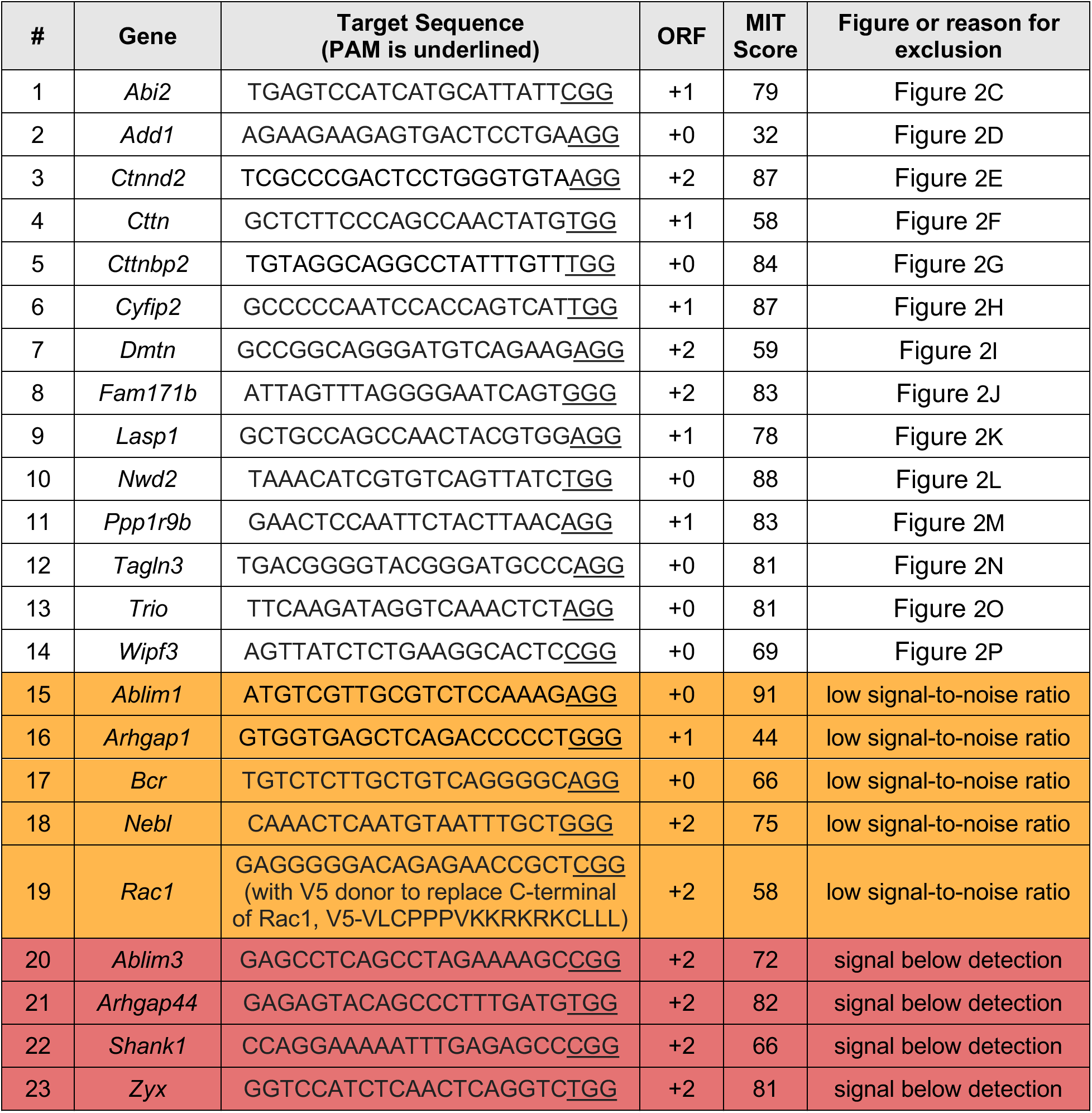
Candidate genes screened for HiUGE validation of the Synapsin-iBioID proteome. A detailed list of C-terminal guide RNAs to validate the presynaptic localization of candidate genes.

